# Conformal Bioprinting of Bi-phasic Jammed Bioinks, Independent of Gravity, Orientation, and Curvature

**DOI:** 10.1101/2025.05.16.654553

**Authors:** Sushant Singh, Lihua Wei, Ehsan Samiei, Khaled Gaber, Qiushi Gao, Aaron H. Persad, Teodor Veres, Axel Günther

## Abstract

Rapid in situ bioprinting on complex, human-scale anatomical surfaces remain a key challenge for clinical translation. Here, we present a gravity-independent, conformal bioprinting strategy using bi-phasic granular bioink and multinozzle printheads capable of adapting to arbitrary surface curvatures. The bioink comprised of jammed gelatin microgels suspended in a fibrinogen matrix exhibits yield-stress behavior to maintain shape fidelity after extrusion while supporting cell viability and proliferation. Two monolithic multinozzle printhead architectures with identical bioink delivery networks were evaluated: (1) a rigid configuration for handheld bioprinting and (2) a soft robotic variant capable of real-time curvature adaptation via pneumatic actuation. Microgravity experiments aboard a parabolic flight confirmed successful bioink deposition under ∼0 g conditions. A ladder-rung microfluidic architecture ensured uniform bioink delivery across printhead nozzles, improving deposition consistency. In situ bioprinting on anatomical facial phantoms confirmed conformal, high-throughput (deposition at 20 mm·s^-1^) deposition of bioink over physiologically relevant curvatures, both with and without cells. Cell-laden constructs retained >85% cell viability post-printing and supported proliferation. This work introduces a scalable bioprinting platform suitable for clinical, remote, and deep-space environments, enabling autonomous tissue fabrication. The curvature-adaptive printhead advances current in situ bioprinting capabilities, facilitating the generation of personalized grafts with complex anatomical geometries.

## 1. Introduction

Human anatomy is defined by inherently three-dimensional structures that span a wide range of curvatures, with radii (*R*) varying from the nanometer to the meter scale (10^-9^ to 10^2^ m).[^1^] These complex surfaces are characterized by two principle curvature radii, *R_1_* and *R_2_*, oriented orthogonally, where typically *R_1_* > *R_2_*. The product of these radii defines the Gaussian curvature of the surface, *κ =* (*R_1_R_2_*)^-1^.

At the nanoscale (*R* ∼ 10^-8^ to 10^-6^ m), curvature plays a pivotal role in biological processes such as the self-assembly of phospholipid bilayers into cell membranes.[^2^] At the tissue level, curvature influences stem cell localization – epidermal stem cells, for instance, preferably reside within the rete ridges of the skin (*R* ∼ 10^-4^ m), [^2–3^] while intestinal stem cells are found in the highly curved crypts of the gut (*R* ∼ 10^-5^ m).^[2, 4]^ Organ-scale curvature is equally diverse: rigid structures such as bone exhibit minimal dynamic deformation (e.g., femur with *R_1_* ∼1 m[^5^] and *R_2_* ∼1.2 × 10^-2^ m[^6^]). Soft tissues dynamically reshape in response to physiological demands – for example, the left ventricle of the heart changes curvature by ∼81% between systole (*R_1_ ∼*1.6 × 10^-2^ m) and diastole (*R ∼*2.8 × 10^-2^ m).[^7^] Similarly, the flexural radii of extremities and digits (*R* of 2-10×10^-3^ m)[^8^][^9^] enable a wide range of motion and fine motor skills. The human face (Fig. 1A) exhibits particularly complex curvature profiles,[^10^] including convex regions like the forehead (*R* ∼ 0.1 m), flatter or concave regions such as the cheeks (*R* ∼ 7-9 × 10^-2^ m), and sharp transitions around the nose and chin (*R* ∼ 1 to 3 × 10^-3^ m), which together support functions ranging from communication and recognition to expression of sensory integration.[^10–11^]

**Figure 1.**
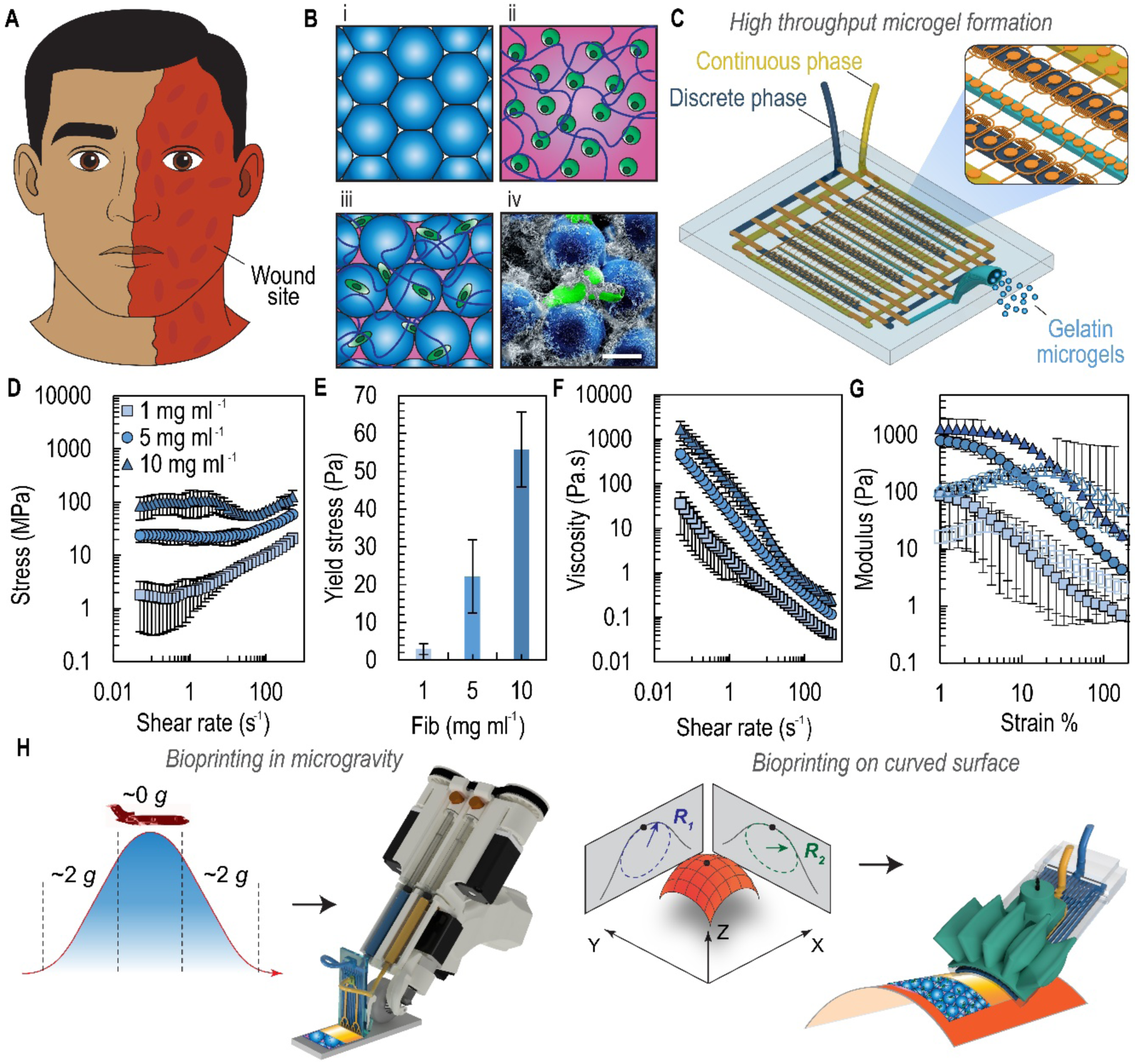
Bi-phasic granular bioinks and soft robotic actuated printhead for bioprinting. (**A**) Schematic of a patient with facial wounds exhibiting various curvatures. The bioprinting process specifically targets convex surfaces such as the forehead, chin, and nose. (**B**) Composition of the bi-phasic polymeric jammed bioink. (**i**) Gelatin microgels are first jammed and then combined with (**ii**) fibrinogen and a cell suspension to form (**iii**) the bi-phasic granular jammed bioink. (**iv**) A scanning electron micrograph of an actual sample highlights cells (in green) within the interstitial spaces of the polymerized bi-phasic bioink Scale bar, 25 µm. (**C**) Schematic of the parallelized droplet microfluidic device containing 102 flow-focusing generators (FFGs) (inset). Rheological characterization of the bioink: (**D**) Stress vs shear rate, (**E**) yield stress, (**F**) viscosity vs shear rate and (**G**) storage (closed symbol) and loss (open symbol) modulus for bi-phasic granular bioinks with varying fibrinogen concentration *C_F_*. (**H**) Bioprinting process using a rigid multinozzle printhead in a microgravity flight and a pneumatically actuated printhead for the co-extrusion of bi-phasic granular bioink and thrombin onto a curved surface characterized by principal curvature radii *R_1_* (parallel to printing direction) and *R_2_* (perpendicular to printing direction).

In-situ bioprinting technologies[^12^] aim to deliver cells and tissue precursors directly onto patient autonomy, by passing the need to manipulate delicate constructs post-fabrication. These approaches have been applied to both external[^13^] and internal[^14^] tissues in clinically relevant environments. Beyond operating rooms, there is growing interest in portable bioprinting systems for use in remote and austere settings, where immediate evacuation may not be feasible.[^15^] For effective use in both clinical and extraterrestrial contexts, in situ bioprinting platforms must meet three critical requirements: (1) Gravitational independence, maintaining structural fidelity in microgravity (0 *g*), lunar gravity (0.17 *g*), Martian gravity (0.38 *g*), and Earth gravity (1 *g =* 9.81 m·s^-2^); (2) Orientation independence, functioning effectively across a full range of angles (0–90°) to reduce the need for patient repositioning; and (3) Geometric adaptability, conforming to anatomies with both positive (convex) and negative (concave) Gaussian curvatures.

To meet these demands, we employ a bi-phasic granular bioink composed of jammed gelatin microgels suspended within a fibrinogen matrix (Fig. 1B).[^16^] Here, the microgels are produced using a parallelized droplet microfluidic device[^27^] with 102 flow-focusing generators (FFGs), as shown in Fig. 1C. The jamming of the microgels imparts yield-stress behavior to the bioink (Fig. 1D-G), rapid structural recovery after deposition and providing superior printability compared to biopolymers commonly used in 3D cell culture and translational research, such as fibrinogen and thrombin alone.[^17^] In our bioprinting approach, a thrombin solution is co-delivered atop the bi-phasic bioink, inducing fibrinogen polymerization. This presents a significant improvement over our previous formulation,[^13a^] where a fibrinogen-hyaluronic acid mixture (without microgels) offered only moderate shear-thinning properties and limited drainage-free deposition to thicknesses approximately one-tenth that of healthy human dermis (∼1mm).[^18^] Beyond enhancing printability, the interstitial spaces between microgels are expected to promote molecular transport and facilitate cell migration. However, to enable clinical and deep-space applications, a compact, single-use, and robust printhead capable of conforming to complex physiological curvatures is also required. Importantly, design rules for uniformly distributing yield-stress fluids[^19^] remain less well established compared to Newtonian and weakly non-Newtonian fluids,[^20^] necessitating additional validation for consistent bioink delivery.

To address these challenges, we evaluate two multinozzle printhead architectures tailored for different applications: (1) a rigid printhead integrated into an updated handheld bioprinter for extraterrestrial and remote use, and (2) a soft robotic actuated printhead for conformal bioprinting on patient-specific, curved surfaces (Fig. 1H). The soft printhead incorporates pneumatically actuated soft robotic elements that enable dynamic curvature adaptation. This allows the multinozzle array to conform to surfaces with non-zero Gaussian curvature – those possessing positive curvature radii along both the deposition, *R*_1_, and the transverse, *R*_2_, directions. Pneumatic actuation allows precise matching to *R*_2_, facilitating real-time surface adaptation during bioprinting. We refer to this approach as Bioprinting Rapidly Using a Shape-Adaptable Head (BRUSH).

Soft robotic systems have previously demonstrated advantages in actuation and control[^21^], locomotion [^22^], object manipulation[^23^] and controlled material transfer,[^24^] owing to their compliance, low contact pressure, and resilience – qualities that are particularly valuable for biological applications. However, existing bioprinting devices leveraging soft robotic elements are so far limited to continuum robots integrated with single nozzle printheads ^[14a, 25]^, which inherently limit deposition speed and area coverage. By contrast, our multinozzle strategy significantly increases bioprinting throughput, enabling rapid coverage of large wound areas. Previous handheld bioprinting systems by our group ^[13a, 26]^ improved area coverage compared to single-nozzle printheads,[^12^] yet lacked adaptability to curvature perpendicular to the printing direction. By integrating a deformable, soft robotic multinozzle printhead, we now enable real-time, omnidirectional curvature adaptation while maintaining uniform bioink delivery.

In this study, we focus on three objectives using bi-phasic granular bioinks: (1) achieving gravity-independent bioink extrusion and structural fidelity under microgravity (∼0 g) conditions; (2) enabling conformal in situ bioprinting on physiologically relevant convex surfaces; and (3) fabricating large cellular structures (>10⁻³ m²) over complex geometries, including human facial topographies. By combining bi-phasic granular bioinks with multinozzle printheads, we introduce a scalable bioprinting platform for regenerative medicine, space exploration, and remote trauma care in remote or austere environments.

## 2. Results and Discussion

### 2.1. Tunable Bi-Phasic Granular Bioinks

Bi-phasic granular microgel bioinks enable precise and systematic tuning of both rheological and biomaterial properties. To produce gelatin microgels, we designed parallelized FFGs, adapted from previous works,[^27–28^] achieving a throughput of ∼16 mL h^-1^ (Fig. S1). This throughput is critical for generating sufficient quantities of gelatin microgels, accounting for material loss (∼ 15%) during washing and crosslinking. For example, processing 100 mL of gelatin solution yields approximately 85 mL of jammed microgels.

Following extraction from the oil phase, the microgels were crosslinked using microbial transglutaminase (mTG), chosen for its low cytotoxicity[^29^] compared to alternatives such as glutaraldehyde. Our crosslinking protocol (Fig. S2), a slightly modification of a previously published method,[^29b^] ensures the stability of microgels under cell culture conditions (Fig. S3). A jammed bioink was prepared by compacting the microgel suspension to a volume fraction exceeding 64%,[^30^] as confirmed by confocal microscopy.[^31^] The void fraction was subsequently adjusted by adding a fibrinogen solution (Fig. S4), with a 1:10 dilution ratio selected for the remainder of the study, yielding a void fraction of 32.7%.

During bioprinting, the granular bioink and thrombin solution were co-delivered from separate syringes. Thrombin diffused into the fibrinogen matrix, inducing rapid clotting and enhancing structural stability (Fig. 1C).

We observed that increasing the fibrinogen concentration, *C_F_*, systematically elevated the yield stress, viscosity, and extended the linear viscoelastic regime of the storage modulus (Figs. 1D-G). This tun tunability allows fine control over rheological parameters while maintaining a constant void fraction. Notably, the Non-Newtonian behavior of the granular bioinks deviated from the classical Herschel-Bulkley model,[^32^] likely due to cohesive interactions between crosslinked gelatin microgels (Fig. 1D).

The bioink also exhibited robust recovery behavior across consecutive strain cycles (Fig. S5), demonstrating desirable yield-stress properties that support high-fidelity bioprinting. To ensure accurate rheological measurements, we employed a parallel-plate rheometer with sandblasted plates to minimize slippage.[^32^] It is important to note that previous rheological assessment of granular bioinks often utilized smooth-walled plates,[^33^] which can introduce measurement artifacts.[^34^] Slippage arises from microgel depletion near the plate surfaces, leading to steep local velocity gradients and an underestimation of the true apparent viscosity.

### 2.2. Uniform Delivery of Granular Bioinks Using Multinozzle Printheads

While granular bioinks have garnered significant interest for their unique mechanical and biological properties[^35^], their behavior under confined flow conditions, particularly within multinozzle printheads, remains incompletely understood. The widely used Herschel-Buckley model fails to accurately predict the flow behavior of granular bioinks when the microgel-to-channel size ratios exceeds 0.01.[^30^] Goyon *et al*.[^30^] addressed this limitation by introducing a generalized fluidity model to describe jammed emulsion flow under confinement – an approach particularly relevant to extrusion bioprinting. Although extensions of this model have been applied to polymeric microgel systems, they have largely focused on high aspect ratio single-channel geometries to simulate parallel plate configurations.[^32^] Consequently, designing multinozzle printheads that achieve uniform distribution of granular bioinks from a common inlet across multiple microchannels requires new fluidic design strategies.

Here, we compared two internal network architectures for multinozzle printheads: a bifurcated tree structure and a ladder-rung layout (Fig. S 6 and 7). The ladder-rung design was based on flow resistances of daughter channels according to Eq. S1. Additionally, informed by the work of Mansard *et al*.[^36^] on confined granular flows, we incorporated wavy wall features (25 µm amplitude, 400 µm period) to improve flow behavior. These patterned walls introduce local perturbations that enhance microgel mobility normal to the flow direction, mitigating otherwise plug-like flow behavior typical for jammed suspensions.

To assess flow uniformity, we imaged the nozzle array during filling and measured the interval (*Δt*) between the filling of the first and the last of the 16 exit channels. Results showed that for bifurcated architectures, rough-walled channels filled more uniformly smooth-walled counterparts. Specifically, rough-walled bifurcated printheads completed filling 3.6× faster than smooth-walled ones (*Δt* = 1.2 s for rough walls), (Fig. S6).

Comparative analysis revealed that the ladder-rung architecture provided superior bioink distribution compared the bifurcated design (Fig. S7). Interestingly, this advantage persisted regardless of wall roughness. We attribute this behavior to the ladder geometry being less influenced by local variations of fluid viscosity in contrast to the bifurcation design, where viscosity differences at each bifurcation level and cross-talk between channels, driven by varying shear stress, may lead to uneven flow distribution.[^37^] Furthermore, the ladder-rung design demonstrated a lower susceptibility to clogging, reinforcing its practicability for bioprinting applications.

Importantly, bioprinted sheets produced with the optimized ladder-rung printheads exhibited consistent void fractions (Fig. S8), confirming the stability and homogeneity of the constructs. Together, these results establish the ladder-rug architecture as a robust strategy for uniform, high-throughput delivery of granular bioinks in multinozzle extrusion bioprinting.

### 2.3. In-situ Bioprinting Independent of Gravity

Parabolic flight experiments were conducted aboard a modified Falcon 20 aircraft[^38^] (Fig. S9). Due to operational constraints and the short microgravity duration, incorporating an onboard pressure source to actuate the soft robotic printhead was impracticable. We therefore used a printhead configured with a shared fluidic architecture for bioink and crosslinker deposition (Fig. 1A), but without the pneumatic actuation layer. The flight trajectory (Fig. 2B) provided approximately 20 s of microgravity during each parabola, offering a unique window for bioprinting. All experimental manoeuvrers were carefully pre-choreographed.

**Figure 2.**
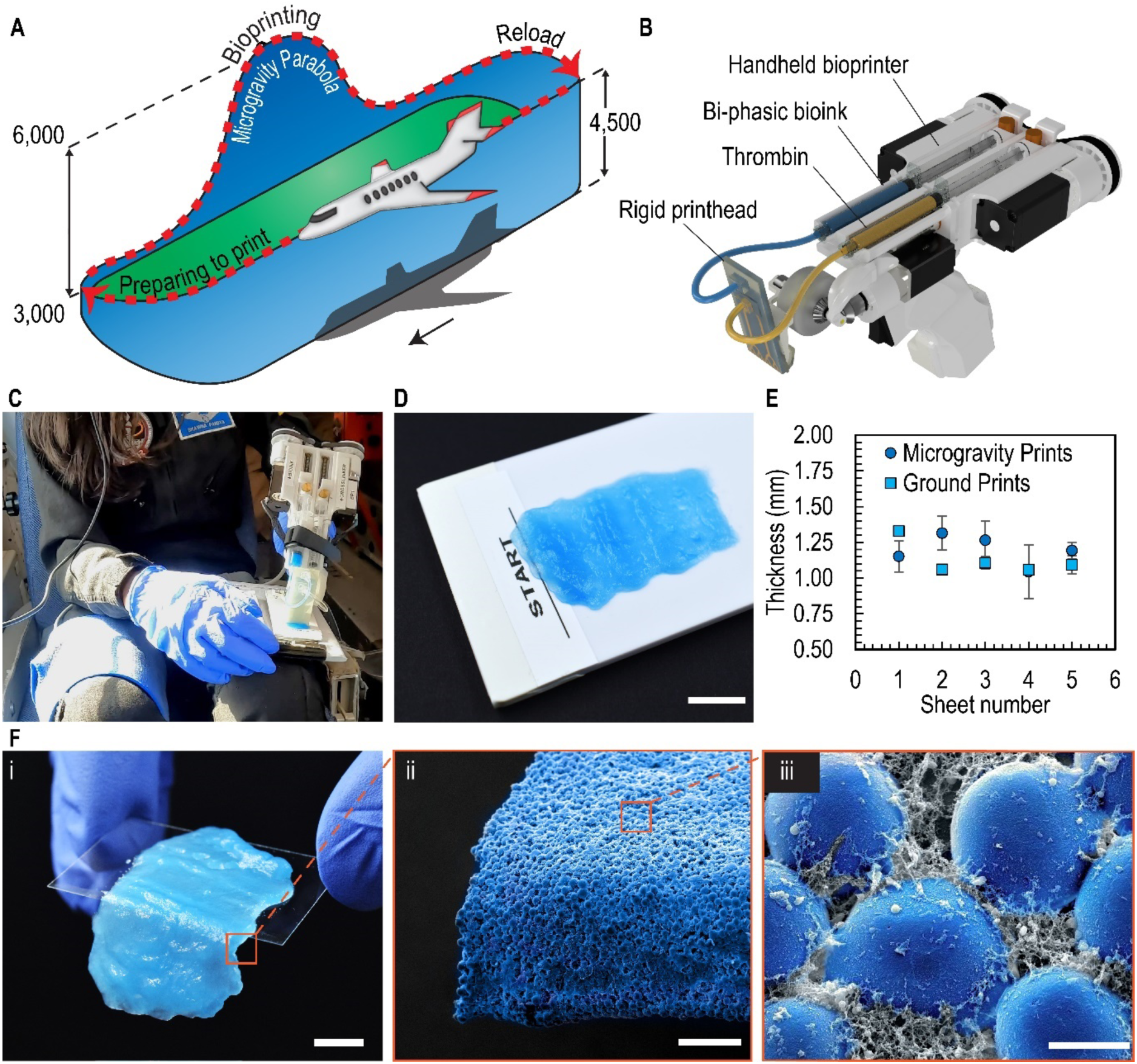
In-situ bioprinting is independent of gravity. (**A**) Flight path for the bioprinting experiment, illustrating the microgravity parabola, which provides approximately 20 s for bioprinting. Two parabolas are performed consecutively using the two onboard bioprinters, followed by a level flight phase that allows for reloading of both bioprinters. All dimensions are in meters. (**B**) Modified handheld bioprinter with integrated multinozzle rigid printhead for delivery of bi-phasic granular bioink and thrombin. (**C**) Operator 1 positions the bioprinters on custom substrates that are secured with Velcro to a leg strap, allowing for easy removal. Upon initiation of the microgravity phase, bioprinting begins and lasts for approximately 10 s. (**D**) As shown, printed sheets adhere to the underlying substrate during microgravity. The sample storage boxes keep the bioprinted sheets hydrated. Samples transported 3 days after bioprinting maintain integrity. (**E**) Three days after printing, the samples are measured using 3D laser scanning. Here, we report thickness data from five different sheets printed on the ground and under microgravity (scale bar, 10 mm). (**F**) Macrographs of the lifted printed constructs confirm structural integrity, while scanning electron micrographs verify fibrinogen clotting, proving that the observed discrete microgels are assembled by the fibrin clot. Scale bars in (i) 10 mm; in (ii) 250 µm; in (iii) 25 µm.

Two operators were involved: Operator 1 conducted bioprinting during microgravity phases, while Operator 2 managed post-print sample handling and storage in a custom-designed box (Fig. S 9 and 10). To minimize dehydration of printed constructs, the storage box was pre-saturated with water, and humidity levels were recorded upon each opening (Fig. S11). An accelerometer attached to the box tracked acceleration profiles during the brief microgravity periods (Fig. S12). The system was operated by both medical and non-medical personnel to assess the influence of different users on bioprinting outcomes (Fig. 2C).

Given the constrained printing window, the bioprinting system was upgraded to achieve extrusion rates approximately 45-fold faster and deposition speeds approximately six-fold higher than in our previous studies (Figs. S13 and S14).^[13a, 26]^ During consecutive microgravity and hypergravity phases (2G), granular bioinks successfully adhered to the substrate (Fig. 2D). Printed sheet dimensions slightly exceeded predictions based on the continuity equation (Fig. 2E), likely due to reduced operator dexterity under weightlessness (as noted by Operator 1) and flight-induced turbulence. The coefficient of variation in sheet thickness during microgravity printing was 8.6%. Ground-based experiments, free from such perturbations, showed improved consistency, with a reduced coefficient of variation of 4.3%.

Operator-dependent variability was also observed, particularly related to the force affecting translation speed, particularly when it exceeded the torque capabilities of the stepper motors. Nevertheless, clotting of 1-mm-thick sheets was successfully achieved, and scanning electron microscopy confirmed fibrin network formation (Fig. 2F).

To our knowledge, this work represents the first demonstration of bi-phasic granular bioink deposition under microgravity conditions. While a handheld bioprinter featuring mechanical synchronization of biomaterial extrusion and printhead translation was recently aboard the International Space Station,[^39^] our approach provides a pathway toward scalable, robust in situ bioprinting in low-gravity environments.

### 2.4. Soft Robotically Actuated Multinozzle Printheads

To enable bioprinting onto curved surfaces, we integrated a pneumatically actuated soft robotic layer atop the microfluidic layers for the delivery of bioink and cross-linker. This design yields a monolithic, curvature-adaptable printhead capable of consistently depositing uniform-thickness bioink layers on convex surfaces. The bottom feature layer, fabricated via soft-lithography, distributes a pressure-driven flow of bi-phasic granular bioink from a single inlet (Fig. S15A), into an inch-wide microchannel array for uniform deposition. Above it, a second soft lithographically patterned layer delivers the thrombin cross-linker and incorporates a liquid metal-based curvature sensor.[^40^]

The electrical resistance of the curvature sensor is determined by the cross sectional area and length of its microchannels,[^41^] which is carefully designed to keep the printhead surface temperature below physiological thresholds during Joule heating, thus preventing cell damage or bioink denaturation. Finally, a soft robotic actuation layer[^42^] is bonded atop the microfluidic stack, enabling the exit section of the multinozzle printhead to bend and match the target curvature radius (*R_2_*) by applying controlled pneumatic pressure (Fig. 3A).

**Figure 3.**
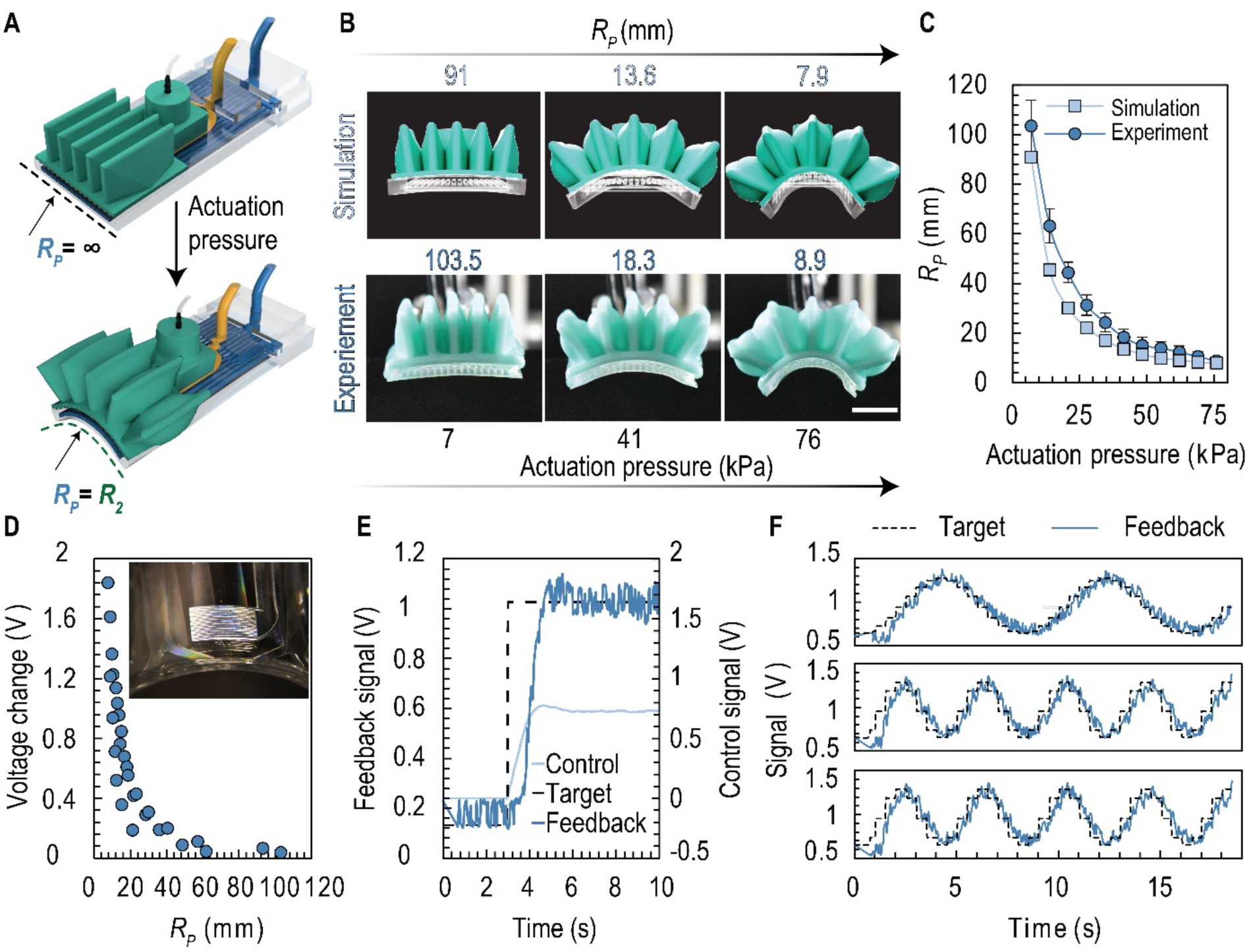
Soft robotically actuated multinozzle heads for bioprinting. (**A**) Simulation and experimental results of printhead deformation from soft robotic actuation (scale bar, 10 mm). (**B**) Curvature radius of printhead (*R_P_*) vs actuation pressure for the printhead during loading phase. Experimental result among three independently made devices is compared to simulation. (**C**) Voltage change recorded from the curvature sensor at different *R_P_* during the loading phase. Datapoints are collected from three independently made devices. (**D**) Printhead reaches the target *R_2_* of 10 mm (converted to voltage signal) from completely flat condition; target was reached in below 1.5 seconds. (**E**) Close-loop control of the printhead with sinusoidal target profile (*R_2_*, converted to voltage signal) from 10 mm to 50 mm at three frequencies (0.125, 0.25 and 0.5 Hz). Sampling frequency for target *R_2_* is 2 Hz in all cases.

The design of the soft robotic layer was informed by finite element simulations to ensure uniform lateral curvature and accommodate a broad range of curvatures (*R_P_*). Experimental measurements different actuation pressures showed favorable agreement with predictions from simulations (Fig. 3A and Fig. S15B), even at pressures beyond the operational range (0 – 76 kPa). Across three different printheads, the standard deviation in curvature measurements remained below 25% during the loading phase (Fig. 3B) and relaxation (Fig. S15C), indicating reliable performance. The actuation layer withstood burst pressures up to 103 kPa (15 PSI) (Fig. S15 D and S15E), demonstrating strong chip-to-world interconnects and robust adhesion, critical for rapid printhead exchange without tubing replacement.

Material selection for the actuation layer ensured operation within the non-hyperelastic regime (Fig. S16), ensuring a near linear curvature change as a function of applied pressure. The curvature sensor is electrically interfaced via a Wheatstone bridge circuit with an operational amplifier and low-pass filter (Fig. S17A). With a 5V input voltage, the printhead surface temperature at sensor location remained below ∼30 °C (Fig. S17B), preserving bioink integrity. During operation, the voltage readout exhibited consistent trends during both loading (Fig. 3C) and relaxation (Fig. S17C) phases across multiple devices, allowing for accurate real-time curvature tracking.

To achieve automated curvature control, a closed-loop feedback system was developed. By adjusting pneumatic pressure in real-time based on the disparity between the measured *R_P_* and the target *R_2_*, the printhead could match a target curvature (*R_2_* = 10 mm) within 1.5 s and maintain it (Fig. 3D). To simulate physiological conditions, we imposed a harmonically varying curvature profile along the deposition direction, updating every 0.5 s. Across different target frequencies (0.125, 0.25 and 0.5 Hz), the feedback system successfully tracked dynamic curvature changes between *R_2_* = 10 mm and 50 mm with minimal phase lag (0.3 s, Fig. 3E), validating the responsiveness of the system for real-time bioprinting on complex surfaces.

### 2.5. In-situ Bioprinting Independent of Curvature and Inclination

We successfully printed different bioink formulations with varying fibrinogen concentrations (*C_F_*) by mounting a multinozzle printhead featuring rough-walled microchannels for lateral distribution of granular bioink to modified 3D printer (Fig. S18). Scanning electron microscopy (SEM) imaging revealed distinct fibrin fiber morphologies formed between the microgels for each formulation (Fig. 4A). Owing to the inherent yield stress behavior of the granular bioink, deposition was achieved on flat, inclined, and curved surfaces (Fig. 4B), with consistent feature fidelity under each condition (Fig. 4C). Anti-fibrinogen staining and SEM analysis at the interface between the granular bioink and the underlying agarose substrate confirmed complete fibrin clotting throughout the constructs (Fig. S19), ensuring strong adhesion.

**Figure 4.**
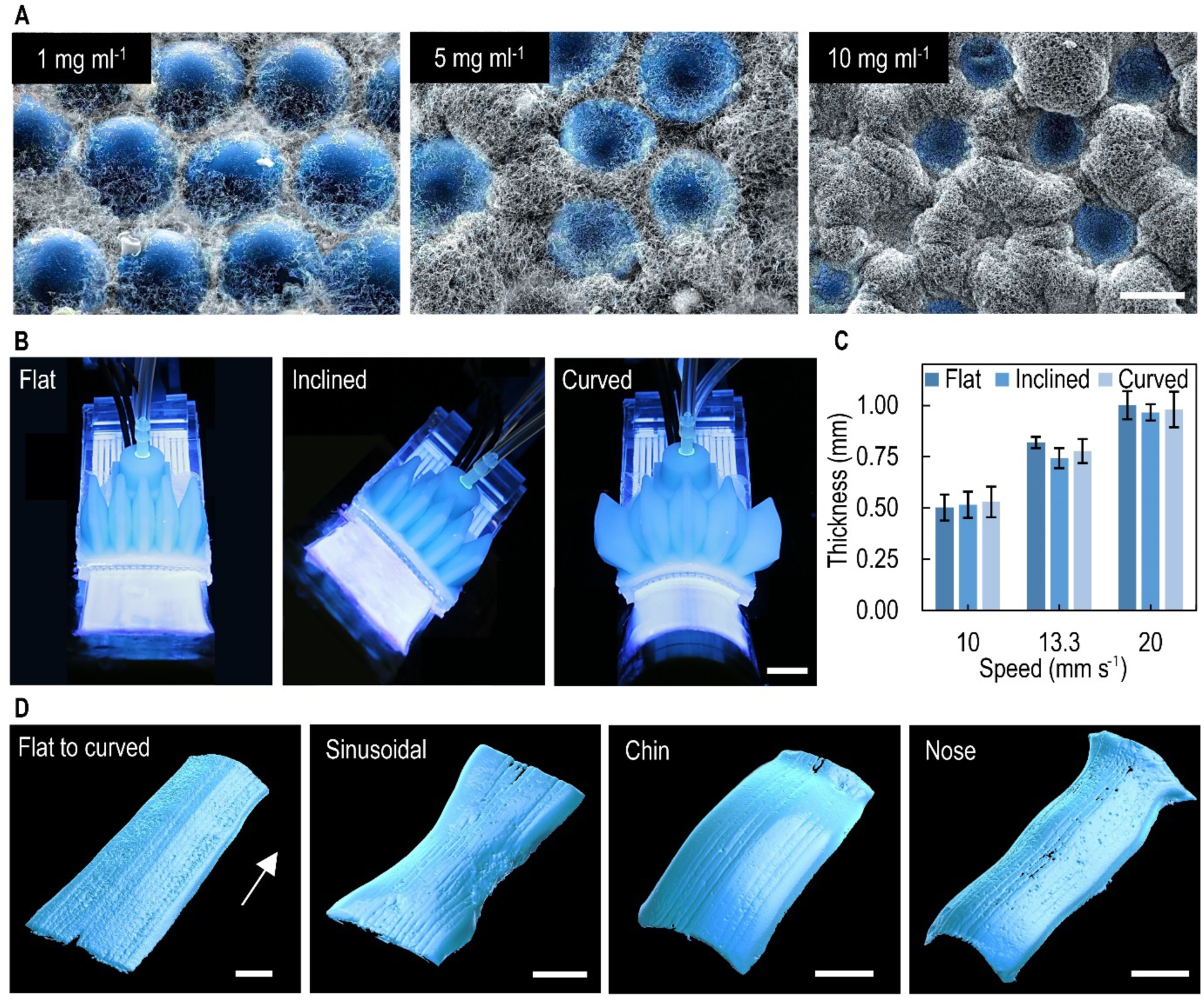
BRUSH-enabled in-situ bioprinting is independent of deposition surface orientation and convex curvature. (**A**) Scanning electron micrographs of in-situ deposited layers prepared with different *C_F_* values show an increase in the relative density of the fibrin clot. Scale bar, 25 µm. (**B**) The BRUSH system is used for bioprinting on flat, inclined (Θ = 45°), and curved (*R_2_* = 25 mm) surfaces using biphasic bioinks and thrombin. The yield-stress behavior of the bioink enables retention of printed features while facilitating clotting of fibrinogen. (**C**) Thickness measurements of bioprinted sheets using 3D laser scanning confirm that target thicknesses of 0.5 mm, 0.75 mm and 1 mm can be achieved. (**D**) Closed-loop bioprinting using BRUSH on substrates with varying curvatures [arrow marks the bioprinting direction]: (i) a linear curvature gradient of 0.02 mm⁻¹ per cm and (ii) a sinusoidal topology where *R_2_* varies from 50 mm at the ends to 10 mm in the center (*R_1_* = 0 in both cases). BRUSH enables in situ bioprinting on physiological surfaces such as chin (*R_1_* ≈ 40 mm and *R_2_* ≈ 15 mm) and nose (*R_1_* ≈ 25 mm and *R_2_* ≈ 8 mm). Scale bar in, 10 mm.

To further evaluate performance, we bioprinted onto engineered surfaces featuring linear and sinusoidal curvature gradients (Fig. 4D), designed to impose significant variations in surface radius (*R_2_*) along the printing direction. These test surfaces were fabricated to maintain a Gaussian curvature of zero, as surfaces remained flat in the direction parallel to the deposition.

A case study highlighted the practical utility of our in-situ bioprinting system: conformal deposition onto anatomically relevant regions of the human face. A 3D scan of a human head phantom (Fig. S20) was used to identify two regions with different *R_2_* values – the chin and the nose (Fig. 4D) representing areas of lower and curvature, respectively. The local mean and Gaussian curvatures matched previously reported values for human facial features[^10^] (Table S1). Molds of the selected areas were fabricated, and the curvature-adaptable printhead dynamically matched the surface curvature during deposition. Successful bioink sheet deposition was validated via micro-computed tomography (micro-CT) image rendering, showing uniform coverage on both anatomical surfaces.

Throughout the deposition, real-time adjustment of the printhead curvature (*R_P_*) along the lateral direction ensured continuous conformity with local variations in *R_2_*. In the extrusion direction, adjustment in the vertical position of the printhead (*z*-axis), coupled with its intrinsic elasticity, accommodated for changes in *R_1_*, maintaining consistent contact between the printhead and the deposition surface.

### 2.6. Bioprinting Cell-Laden Constructs on Physiological Surfaces

To assess the ability of our approach to create large, physiologically relevant cellular structures, we bioprinted human dermal fibroblasts using the bi-phasic granular bioink (Fig. 5A). Post-bioprinting viability remained high, exceeding 85% on both Day 1 and Day 3 (Figs. 5B and 5C), confirming the cytocompatibility of the method. To evaluate the influence of the fibrinogen concentration (*C_F_*) on cell proliferation, EdU staining was performed. Quantification of EdU-positive (EdU+) relative to total nuclei (DAPI+) revealed only a weak correlation with varying *C_F_* values (Figs. 5 D and 4E), suggesting that rheological tuning of the bioink can be achieved without significantly impacting cell proliferation.

**Figure 5.**
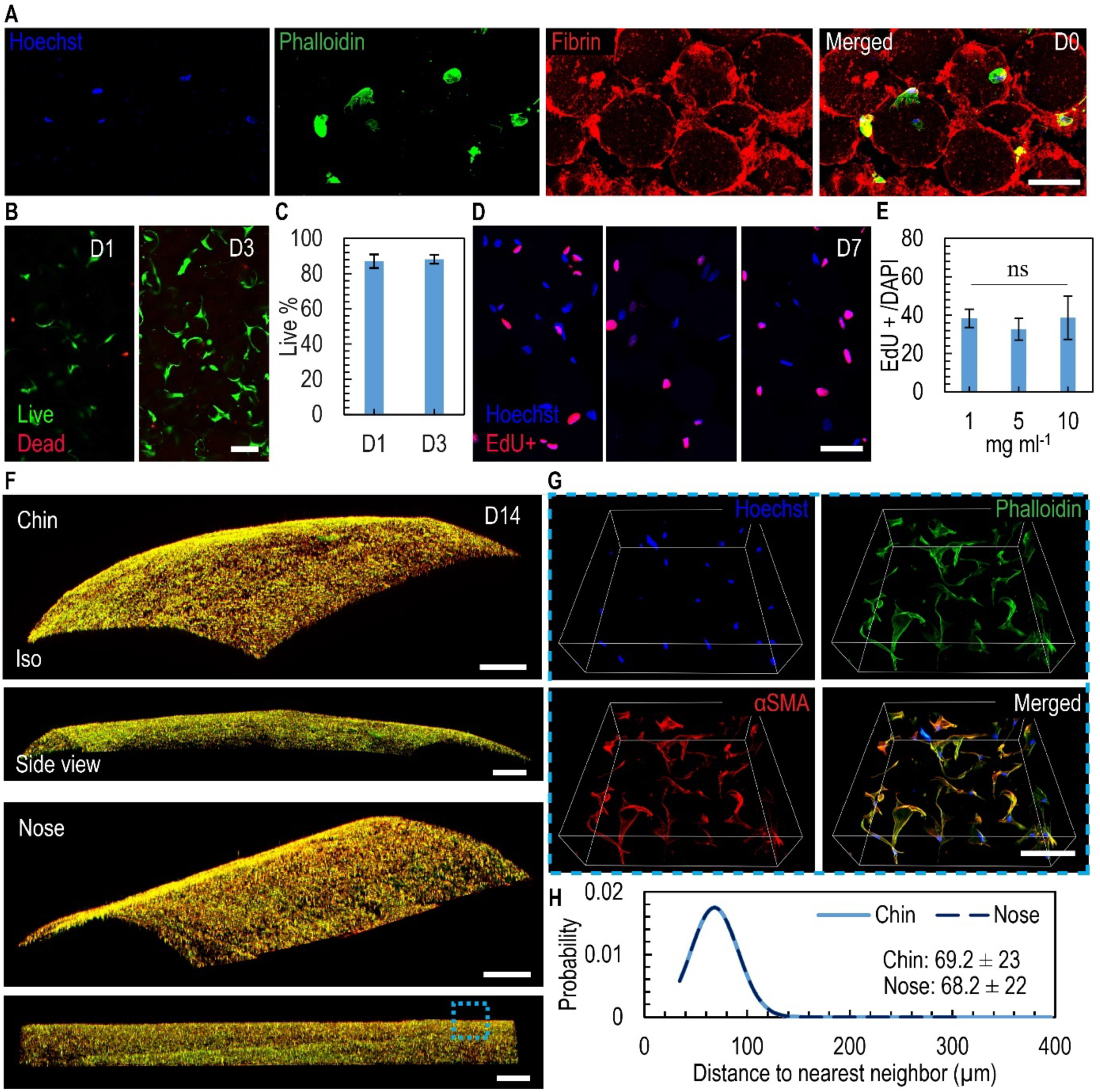
BRUSH-enabled in-situ bioprinting of cellular constructs. (**A**) In-situ deposited layer from cell-laden bi-phasic granular bioink on Day 0 shows delivered cells stained with Hoechst (blue) and Phalloidin (green), reside within interstitial spaces between microgels occupied by fibrin clot (red). Scale bar, 100 µm. (**B** and **C**) Viability of dermal fibroblasts in bioprinted layers after Days 1 and 3 confirms cell survival (n = 5, scale bar, 100 µm). (**D** and **E**) Proliferation analysis using EdU staining of constructs prepared for different values of *C_F_* reveals a non-significant change in proliferation capacity (*n* = 3, scale bar, 50 µm). (**F**) Bi-phasic bioink and dermal fibroblasts are in situ bioprinted onto surfaces that resemble the physiological topologies of the human face, the chin (top) and the nose (bottom) and cultured for 2weeks. Conformally deposited constructs adhere to underlying substrate owing to fibrin’s adhesive properties and promote cell proliferation. Scale bar, 20 mm. (**G**) Dermal fibroblasts occupy fibrin-filled space and organize themselves in three dimensions. Scale bar, 100 µm. (**H**) Distance between nearest neighbor or cell-cell distance remains consistent between samples.

For proof-of-concept validation on curved surfaces, molds from chin and nose regions of the human phantom were employed (Figs. 5F), with an enclosed culture chamber designed to maintain hydration and culture-medium supply during the two-week submerged culture period. Fibrinogen, a clinically approved adhesive, facilitated the stable attachment of the bioprinted sheets throughout culture. After the culture period, the bioprinted constructs exhibited uniform cell coverage without observable contraction or delamination from the substrate. High-magnification imaging (inset, Fig. 5G) revealed that cells were three-dimensionally organized within the fibrin matrix and among the microgel components.

To evaluate the stability of the bioprinted cellular architecture, nearest-neighbor cell distances were measured and found to be consistent across samples (Fig. 5H), confirming spatial uniformity of cell distribution. These results demonstrate the ability of our system to rapidly deposit bi-phasic bioinks onto convex surfaces, achieving effective delivery and retention of cellular payloads.

## 3. Conclusion

While significant advances have been made in bioprinting, major challenges persist in achieving rapid, in situ deposition of bioinks onto complex, human-scale, and irregularly shaped surfaces. In this study, we introduce a scalable bioprinting platform capable of depositing bi-phasic bioinks with high structural fidelity and cell viability, both under normal and microgravity conditions. We report the first multinozzle, shape-deforming printhead featuring monolithically integrated bioink and cross-linker delivery, real-time curvature sensing, and pneumatic actuation. This approach enables gravity-independent, orientation-adaptive, and curvature-conforming bioprinting, overcoming limitations in traditional bioprinting that rely on pre-formed scaffolds.[^43^]

Beyond adaptable printheads, effective in situ bioprinting increasingly depens on personalized surface data for optimizing tissue reconstruction. Advances in 3D scanning technologies have accelerated biomedical applications in implant design,[^44^] wearable customization,[^45^] and anatomical modeling.[^46^] Personalized surface replication across scales^[14b, 47]^ is critical, as implantable tissues and devices must match local curvature, which varies by sex, genetic background,[^48^] age,[^49^] and health status,[^50^] to promote integration[^51^] and improve prosthetic attachment and function.[^52^] Future research should focus on developing next-generation shape-morphing printheads that eliminate the need for pre-scanned surfaces,^[13b, 47c]^ integrate with handheld devices or robotic systems, and employ advanced soft robotic actuation compatible with current clinical workflows. These advances will allow for greater adaptability to complex topologies,[^47c^] including independent controlof principal curvature radii and bioprinting on concave surfaces, representing a critical area for development.

Importantly, in vivo preclinical validation of these adaptable systems will be essential for translating this technology into therapeutic applications.

In summary, this study demonstrates: (1) the use of bi-phasic bioinks to maintain structural fidelity post-extrusion, including under microgravity; (2) conformal bioprinting of cell-laden bi-phasic bioink layers onto physiologically relevant convex surfaces, and (3) the scalability of the platform for fabricating large, cellular constructs. Altogether, this work establishes the foundation for the next generation of customizable, high-throughput, clinically relevant in situ bioprinting systems, with potential applications spanning hospital settings, remote trauma care, and space-based healthcare.

## 4. Experimental Section

### 4.1 Bioink Flow Synthesis

Crosslinked gelatin microgels, the primary component of the bioink, were synthesized using a parallel droplet microfluidic system. A 10% gelatin solution (G9391, Sigma Aldrich, Oakville, ON, Canada) was emulsified in heavy mineral oil (AC415080025, Thermo Scientific, Canada) containing 2% Span 80 (8.40123, Sigma Aldrich, Oakville, ON, Canada) via flow-focusing generators. Flow rates of the oil and gelatin phases were regulated using pressure controllers (T2000 and T3000, Marsh Bellofram, VW, USA) and sensors (SSI Technologies, LLC, WI, USA) connected to a pressurized air line. To prevent premature gelatin gelation, reservoirs and tubing were maintained at 37°C using a water bath and heating belts, respectively. Post-emulsification, to avoid the coalescence of emerging gelatin microgels, the outlet tubing and collection bottle were submerged in a 4-8°C water bath. Microfluidic device fabrication followed previously described protocols.[^28c^]

### 4.2 Bioink Preparation and Rheology Analysis

As previously reported,[^28c^] gelatin microgels were centrifuged at 3000×g for 2 min and diluted in fibrinogen (F8630, Sigma Aldrich, Canada) solutions at varying concentrations. Fibrinogen stock solutions were prepared in phosphate-buffered saline (FBS) unless otherwise noted. Precise control over the fibrinogen concentration (*C_F_*) was achieved by considering the dilution factor.

### 4.3 Handheld Bioprinter

The handheld bioprinter simultaneously extruded bioink (28 mL·min⁻¹) and thrombin (250 U·mL⁻¹, 1.4 mL·min⁻¹, T4648, Sigma Aldrich, Canada) while translating at 1.9 cm·s⁻¹. Dual-syringe extrusion (3, 5, or 10 mL syringes) was achieved using two independently controlled stepper motors (NEMA 11 Bipolar Stepper 11HS20-0674S-PG5, Stepper Online, UK) with integrated gearboxes. A custom silicone wheel (Ecoflex 00-30, Smooth-On, PA, USA; 36.5 mm diameter) driven by a NEMA 8 stepper motor (Bipolar Stepper 8HS15-0604S-PG19, Stepper Online, UK) provided translational motion. An Arduino-based control box synchronized motor control and extrusion timing, triggered via a manual switch for user safety. Stepper motor accuracy was calibrated and verified via rotary encoders (E38S6G5-600B-G24N, Amazon, Canada) attached to bioink and crosslinker stepper motors.

The printhead mimicked the internal microchannel architecture of the soft robotically actuated printhead but was rigidly 3D printed in acrylic resin part with paraffin wax support (multiJet 3D printer ProJet 3500HD Max, 3D System, USA). Wax removal involved oven heating at 65°C, followed by oil submersion (mineral oil, 330779, Sigma Aldrich, Canada) at the same temperature, manual flushing using a syringe, soap rinsing, and ultrasonic cleaning. Tubing (1/16’’ ID, ∼0.1m length, Tygon, VWR International, Radnor, PA, USA) was connected to the bioprinter syringes.

### 4.4 Bioprinting System

The automated bioprinting setup consisted of a modified PRUSA MK2 3D printer (Prusa Research, Prague, Czech Republic) equipped with a soft robotic printhead via a custom connector. Bioink and crosslinker were delivered through 1/16’’ ID Tygon tubing, using syringe pumps (Model PhD 2000, Harvard Apparatus, MA, USA). The soft robotic actuator was driven by either a pneumatic pressure controller and sensor (for deposition) or a manual syringe pump (for curvature sensor characterization). Solid wires linked the embedded curvature sensors to an external measurement system (McMaster-Carr, IL, USA). Printing substrates included either molded agarose blocks or 3D-printed molds, secured with custom acrylic holders (Mcmaster-Carr, IL, USA). Motion control was achieved via a custom G-code executed in Simplify3D, while flow rate and actuation were managed independently. Curvature feedback was achieved through a custom MATLAB script (MathWorks, Natick, MA, USA). The bioprinting flow rates and translational speeds matched those of the handheld bioprinter.

### 4.5 Soft Robotically Actuated Multinozzle Printhead

The printhead contained distinct layers for bioink and crosslinker flows, along with integrated curvature sensing. Daughter channels (height: 1000 µm) and distribution channels (height: 2500 µm) in the bioink layer featured rough wall patterns (12.5 µm amplitude, 400 µm period). Fabrication involved photolithography and soft lithography, as described earlier. Polydimethylsiloxane (PDMS) precursor (Sylgard 184, Dow Corning, Midland, MI, USA) was spread on the master mold, degassed, and an acrylic mold was positioned to control layer thickness. Two half-layers were cured and bonded after corona treatment, ensuring careful channel alignment. The crosslinker and sensor layer was fabricated similarly, with a fluidic channel height of 150 µm and a sensor channel pattern height of 25 µm. A 6 mm thick PDMS slap was prepared for world-to-chip fluidic connections. Inlet holes were punched. The multilayered device was cured in an oven at 80°C for 1 h. Liquid GaInSn alloy (McMaster-Carr, IL, USA) was injected into the sensor channels, and the soft robotic actuator was bonded on top of the printhead.

### 4.6 Soft Robotic Actuator

Rigid male and female molds were designed in SolidWorks (Dassault Systèmes, France) and 3D printed (ProJet MJP 3600, 3D Systems, USA). The actuator was molded using either Mold Star 15 or Ecoflex 00-30 precursors (Smooth-On, PA, USA), mixed 1:1 by mass, degassed, and cast into the female mold. The male mold was inserted into the female mold, aligned, clamped, and cured at 55°C for at least 4 h. The actuator was removed from the mold, and a hole was punched at the desired location to attach a 1-mm ID tube. A 3D-printed rigid ring was inserted at the pressure connection port to prevent overexpansion during pressurization. The actuator was bonded to the printhead using Sil-Poxy adhesive (Smooth-On, PA, USA) and cured at 55°C for an additional 30 min.

### 4.7 Bioink Channel Filling Characterization

Jammed bioink was injected into the printhead’s bioink channels using syringe pumps (Model PhD 2000, Harvard Apparatus, MA, USA). The filling process was recorded at 100 Hz using a high-speed camera (model 1200hs, PCO Imaging, Kelheim, Germany). The channel filling time was quantified by subtracting the timestamp corresponding to the filling of the first daughter channel from that of each subsequent channel.

### 4.8 Tissue Profile Scanning

Printed constructs were characterized by using a laser scanner (FaroArm Quantum, Lake Mary, FL, USA). Agarose substrates were scanned before both pre- and post-deposition. Scan datasets were aligned and compared using Geomagic Control X software (Hexagon AB, Stockholm, Sweden) to measure bioink deposition thickness by overlapping the scans while keeping the pre-print substrate as the reference. To enhance scan contrast, both agarose substrates and bioinks were supplemented with food dyes. The PRUSA 3D printer was immobilized during the scanning process to prevent misalignment. Scanned profiles were subsequently sectioned using the program Magics (Materialise, Belgium) and imported into AutoCAD (Autodesk, San Franscisco, CA, USA) for thickness analysis.

### 4.9 Deposition on Facial Feature Topologies

An intubation head was rented from the Surgical Skills Centre at University of Toronto. The facial structure was scanned via laser scanning, and the topology was reconstructed into CAD models using Magics and Solidworks (Dassault Systemes, Vélizy-Villacoublay, France). Custom molds for agarose substrates and tissue culture devices were designed accordingly. Agarose molds were prepared by dissolving agarose powder in DI water (3% w/v) at 80°C, pouring into 3D-printed molds, and curing at room temperature for at least 15 min. Agarose substrates were positioned in custom acrylic holders mounted on the PRUSA 3D printer for deposition experiments. For tissue culture devices, porous nylon membranes (Tisch Scientific, OH, USA) were attached at the printed facial feature sites using silicone rubber sealant (Momentive, NY, USA).

### 4.10 Cell Culture

Human dermal fibroblasts from adult skin (NHDF-Ad, CC-2511, Lonza, Basel, Switzerland) were cultured in high glucose medium (DMEM, 11965092, Gibco, Thermo Fisher, Canada) supplemented with 10% fetal bovine serum (A5670701, Gibco, Thermo Fisher, Canada) and 1% Penicillin-Streptomycin (15140148, Gibco, Thermo Fisher, Canada). Cells at passages 2–5 were expanded in T75 or T175 flasks, with media exchanges every two days. Cells were harvested by trypsinization and upon reaching ∼85% confluency.

### 4.11 Tissue Culture of Bioprinted Sheets

Cellular bioinks were prepared by suspending harvested fibroblasts in fibrinogen solutions following centrifugation at 300 G for 5 min and media removal. To minimize premature clotting, cells were resuspended in fibrinogen prepared in high-glucose DMEM without fetal bovine serum (FBS). Bioprinting was performed inside a biosafety cabinet onto 2% agarose slabs (3 × 1 × 20 cm) to facilitate handling. Printed constructs were allowed to clot for ∼1 min before transferring into 6-well plates for culture. Cell culture media was replaced every two days.

### 4.12 Immunostaining

***Live/Dead Viability Analysis.*** Samples (1 cm²) were stained using a cell viability imaging kit (R37601, Thermo Scientific, Canada) and incubated on an orbital shaker at (100 rpm) for at least 30 min. Imaging was conducted with a light sheet microscope (Stellaris 5, Leica Microsystems, Wetzlar, Germany) and analysis was performed using Imaris 10.0 (Oxford Instruments, Abingdon, UK).

***EdU Proliferation Assay.*** Samples were incubated overnight with 10 µM EdU (ab222421, Abcam, Ontario, Canada), fixed with 4% paraformaldehyde (PFA) for 15 min, and cryosectioned at 20 µm thickness. EdU detection was performed according to the manufacturer’s protocol, and samples were mounted with ProLong Gold Antifade Mountant (P36930, Thermo Scientific, Canada) prior to imaging.

***Fibrinogen Immunostaining.*** Samples were fixed in 4% PFA, cryosectioned, and immunostained using an anti-fibrinogen antibody (ab27913, Abcam, Canada) per the manufacturer’s instructions.

***Cytoskeletal and Nuclear Staining on Curved Wells.*** Samples from curved well structures were fixed with 4% PFA for 20 min, stained with Phalloidin-FITC (ab235137, Abcam, Canada) for F-actin visualization, and counterstained with Hoechst 3342 (ab228551, Abcam, Canada) for nuclei detection according to the manufacturer’s protocols.

*Statistical analysis:* The data in the graphs are presented as mean +/- standard deviation. Results are based on 3 or more independent experiments. Student t test and one way ANOVA were used where applicable.

## Supporting Information

Supporting Information is available from the Wiley Online Library or from the author.

## Acknowledgements

We thank Dr. Dan Voicu (University of Toronto, UofT) for his support throughout the various stages of microfluidic device preparation and acknowledge insightful discussions with Dr. Keith Morton (National Research Council). We also appreciate Dr. Durgesh Kavishvar from Dr. Arun Ramchandran’s laboratory (UofT) for his advice on setting up the rheology experiments. We extend our gratitude to Dr. Joseph Umoh from the Preclinical Imaging Research Centre at Robarts Research Institute for conducting and preparing raw data for micro-CT scans. We thank Wei Lin for the initial design and proof-of-concept characterization of the liquid metal-based curvature sensor. We are grateful to the International Institute for Astronautical Sciences (IIAS) for assisting with payload integration onto the Falcon 20 aircraft and for in-kind support funding the flight campaign. Special thanks to Dr. Norah Patten and Dr. Shawna Pandya (IIAS) for performing bioprinting experiments in microgravity and helping establish the protocol for in-situ bioprinting aboard the Falcon 20 aircraft.

We acknowledge support from NSERC (AG, RGPIN-2017-06781, RGPIN-2024-06528, I2IPJ 576571-22), the Canadian Space Agency (AG, FAST grant, A 23FATORA3), postdoctoral fellowships (ES, from NSERC, PDF-532965-2019; and the Center for Research and Applications in Fluidic Technologies, CRAFT), and graduate fellowships (SS: Barbara and Frank Milligan Graduate Fellowship; and LW: Ontario Graduate Scholarship and NSERC PGS-D). Device fabrication was conducted at the CRAFT Device and Tissue Foundries, open research facilities supported by the University of Toronto, NRC (Disruptive Technology Solutions for Cell and Gene Therapy Challenge Program), the Canada Foundation for Innovation, and the Ontario Research Fund (Ontario-Québec Center for Organ-on-a-Chip Engineering, Center for Advancing Neurotechnological Innovation to Application).

## Author Contributions

S. Singh and L. Wei were responsible for conceptualization, investigation, data curation, formal analysis, visualization, original draft writing, reviewing and editing E. Samiei conducted preliminary conceptualization, investigation, and validation of the soft-robotic actuators, bioprinting experiments, and droplet generator design and contributed to manuscript review. S. Singh fabricated all droplet generators from the preliminary stage to final implementation, with assistance from K. Gaber for the parabolic flight campaign. S. Singh and K. Gaber prepared the bioprinter payload for the microgravity experiment, conducted validation, and performed analysis under the supervision of A. Persad. L. Wei was responsible for conceptualizing, investigating, and modifying closed-loop controlled printheads. Q. Gao conducted preliminary investigations on the flow of jammed microgels within microfluidic printheads with bifurcations. T. Veres co-supervised students, reviewed and edited the final manuscript draft. A. Günther secured funding, supervised the project, contributed to conceptualization, investigation of all aspects of the project, assisted with the original manuscript draft, and reviewed and edited the manuscript document.

