## Supplementary Information for "Conformal Bioprinting of Bi-phasic Jammed Bioinks, Independent of Gravity, Orientation, and Curvature"


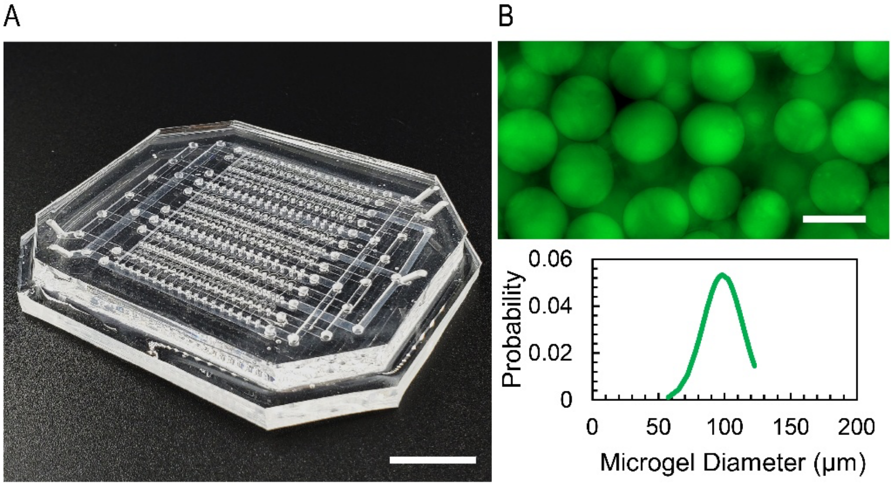


**Figure S1.** Fabrication and characterization of gelatin microgels. (**A**) Photograph of the multilayer soft lithographically patterned device, composed of three polydimethylsiloxane layers (scale bar, 5 mm). (**B**) Fluorescence micrograph of gelatin microgels (autofluorescent at 488 nm) following washing (scale bar, 100 μm) and associate diameter distribution of prepared microgels, determined from measurements over more than 100 microgels.

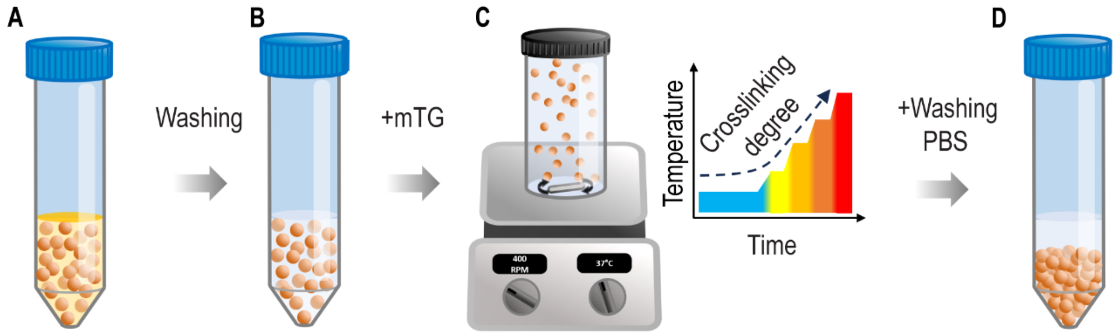


**Figure S2.** Workflow for microgel preparation and transfer. (**A**) Gelatin microgels suspended in mineral oil. (**B**). Following surfactant addition, centrifugation and oil phase removal, microgels are transferred to deionized water (**C**) Microgels are crosslinked with microbial transglutaminase on a hotplate with the temperature increased by 5°C per hour, from room temperature to 40°C. (**D**) Final washing of crosslinked microgels in phosphate-buffered saline (PBS).

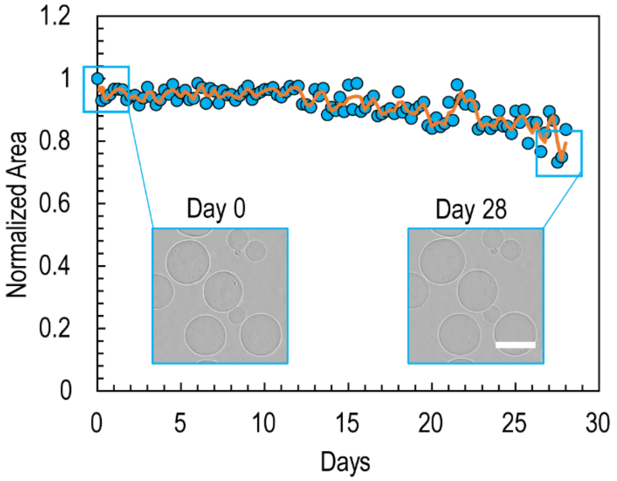


**Figure S3.** Stability assessment of microgels incubated in PBS at 37°C for 28 days. Changes in projected area were quantified from bright-field micrographs, revealing a moderate decrease after 2 weeks (scale bar: 100 μm). Data represents measurements for more than 100 microgels.

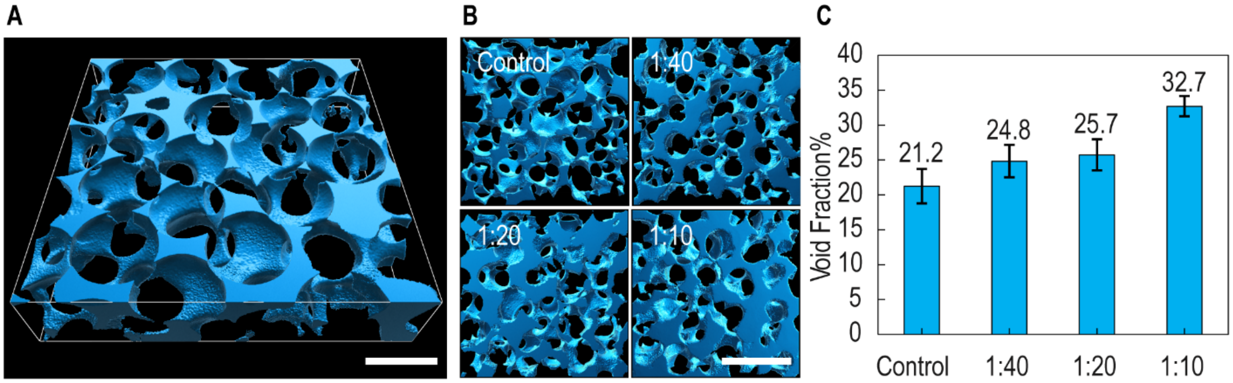


**Figure S4.** (**A**) Three-dimensional rendering of microgel (disperse phase) and fluorescently labelled continuous liquid phase (turquoise) obtained from a confocal z-stack (scale bar: 100 µm), visualized using Imaris 10.2 software. (**B**) Rendered phase distributions for bioinks prepared at different dilution ratios (scale bar: 200 µm). (**C**) Measured liquid fractions corresponding to each dilution ratio.

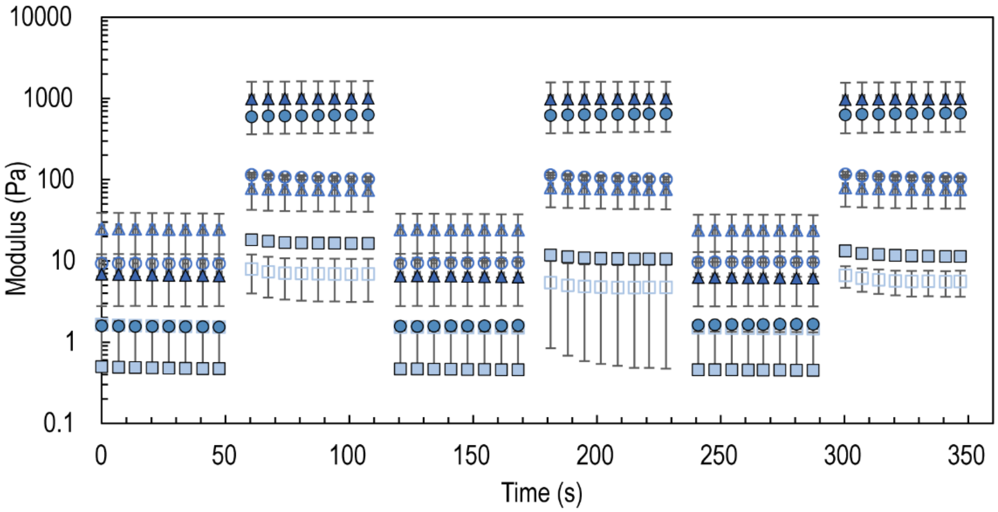


**Figure S5.** Oscillatory sweeps show the recovery response of the bioinks for varying fibrinogen concentrations. Storage modulus reported in solid symbols and loss modulus in open symbols.

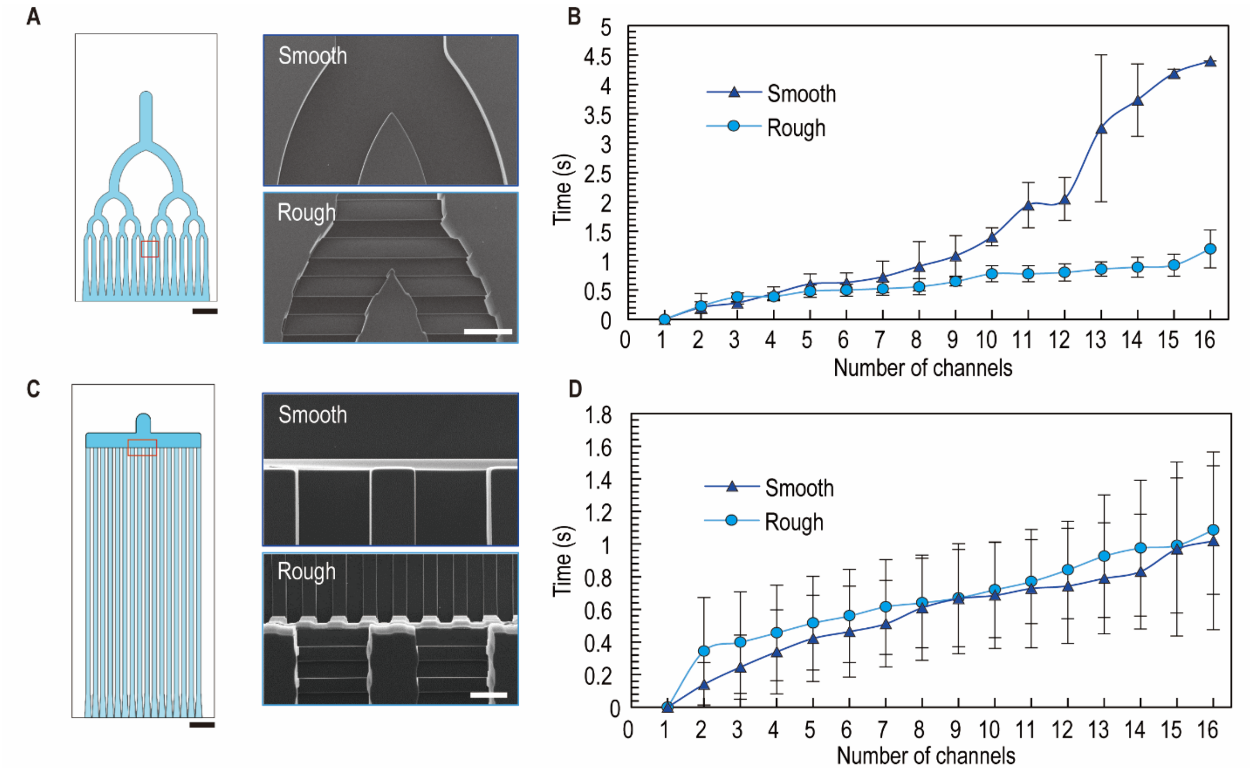


**Figure S6.** (**A**) Representative images showing channel filling at different time points for smooth- and rough-walled bifurcation geometries. (**B**) Quantification of filling time for each daughter channel in bifurcated geometries; filling time is defined as the difference between the time each channel fills and the time the first channel. (**C**) Images of smooth and rough-walled ladder printhead geometries (scale bar: 50 μm). (**D**) Channel filling time measured for the ladder geometries.

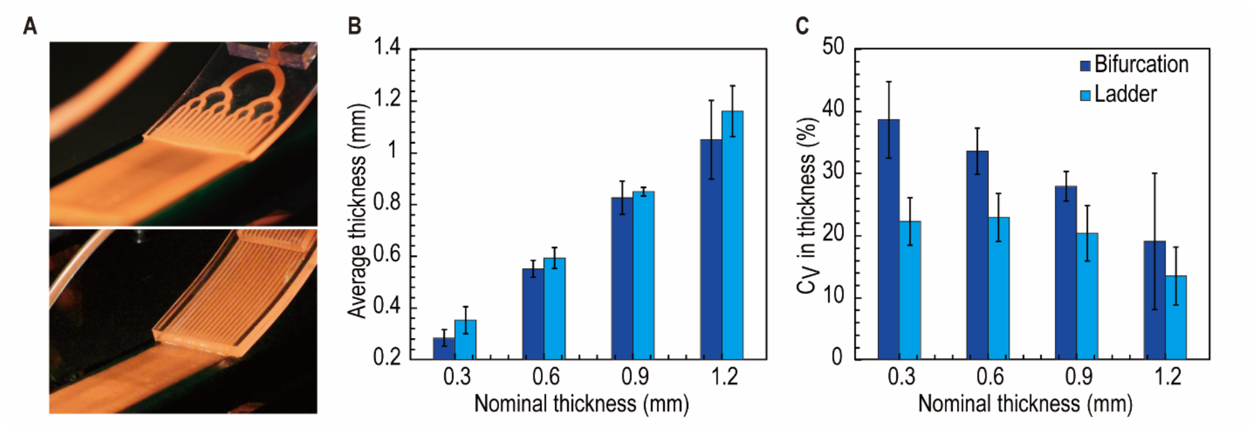


**Figure S7.** (**A**) Photograph of bi-phasic bioink deposition onto flat surface using a multinozzle microfluidic printhead (without cross-linker and soft robotic actuation layers), featuring either bifurcation (top) or ladder (bottom) channel networks, both with roughened side walls. (**B**) Average sheet thickness and (**C**) coefficient of variation in thickness for four different nominal sheet thicknesses, comparing bioink deposition using the two channel network designs.

**Equation S1.** Design rule used for dimensions of ladder geometry. (*n*: number of daughter channels, *L*: channel length, *w*: channel width, *h*: channel height ($h < w$), subscript *p*: dimensions for parent channel, subscript *d*: dimensions for daughter channels)

$$n = \frac{L_d w_p h_p^3}{200 L_p w_d h_d^3}$$

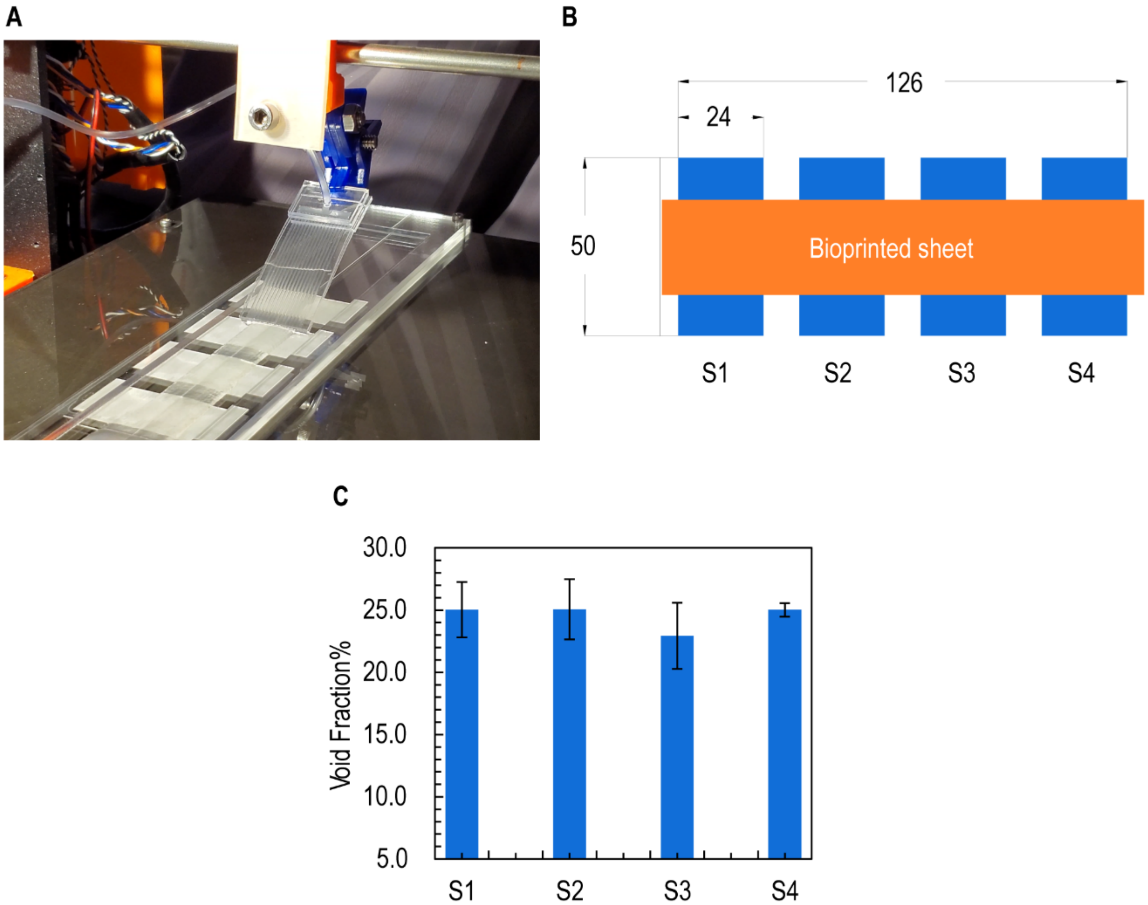


**Figure S8.** Characterization of bi-phasic bioink deposition onto discrete surfaces. (**A**) Photograph of bi-phasic bioink deposition onto glass cover slides arranged with uniform 5 mm spacing. The deposition was performed using soft lithographically patterned multinozzle printhead comprising only the bioink distribution layer, without cross-linker and soft robotic actuation layers. (**B**) Schematic of the deposition setup showing the arrangement and dimensions of the cover slides (all dimensions in mm). (**C**) Quantification of void fraction during deposition, demonstrating consistent print properties.

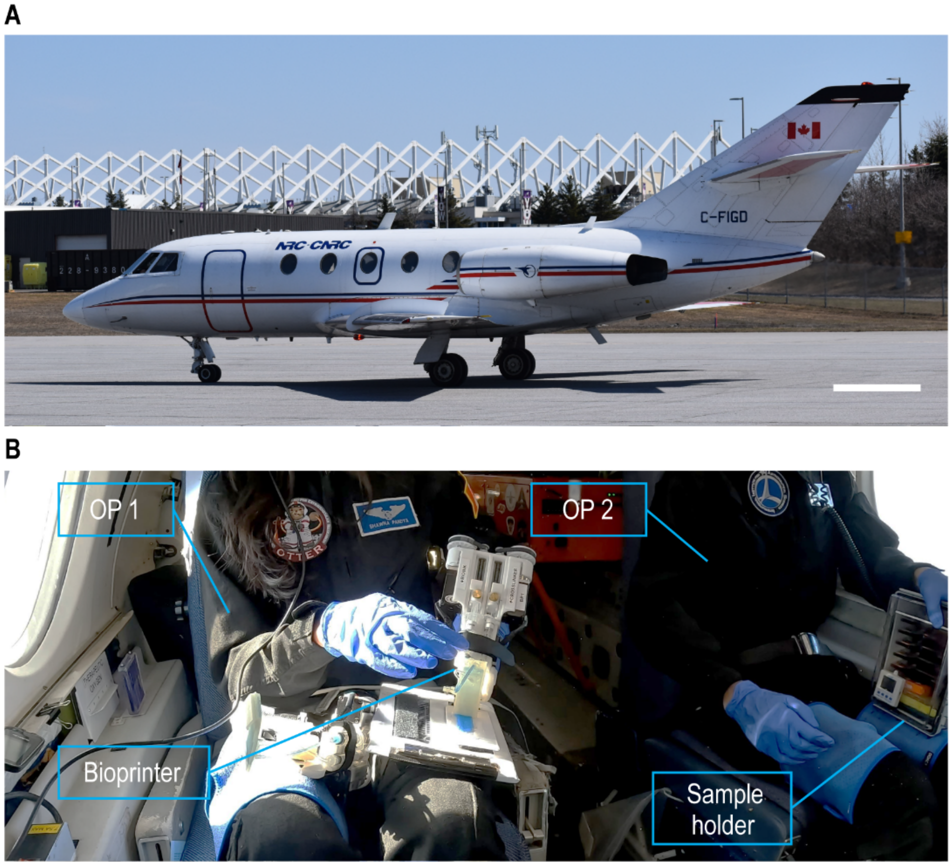


**Figure S9.** Microgravity printing experiment. (**A**) Photograph of the experimental setup aboard a modified Falcon 20 aircraft during parabolic flight (scale bar, 2 m). A rigid version of the multi-nozzle printhead, incorporating both bioink and crosslinker distribution layers, was used for deposition. (**B**) Operational workflow: Operator 1 (OP1) performed bioprinting onto a flat deposition surface secured to their upper leg and subsequently transferred the printed bioink sheet to Operator 2 (OP2), for placement into a custom saturated humidity storage container.

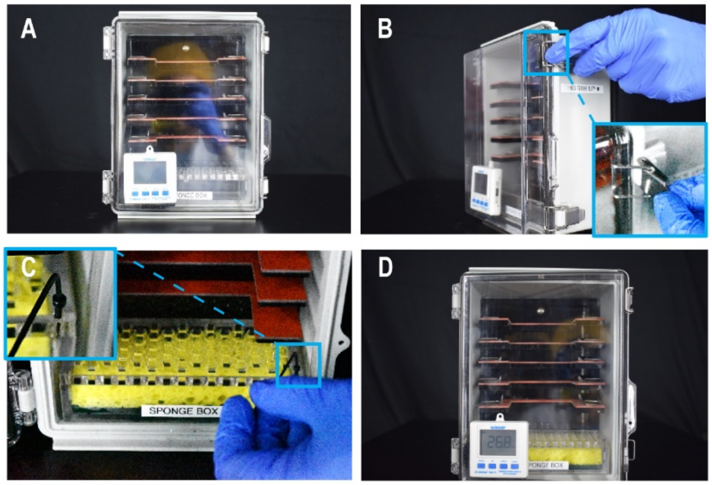


**Figure S10.** (**A**) and (**B**) The sample storage box is designed to maintain the hydration of printed samples. (**C**) A wet sponge is placed inside the box to prevent water movement and maintain saturated humidity. (**D**) Temperature and relative humidity sensors continuously monitor the environmental conditions within the box.

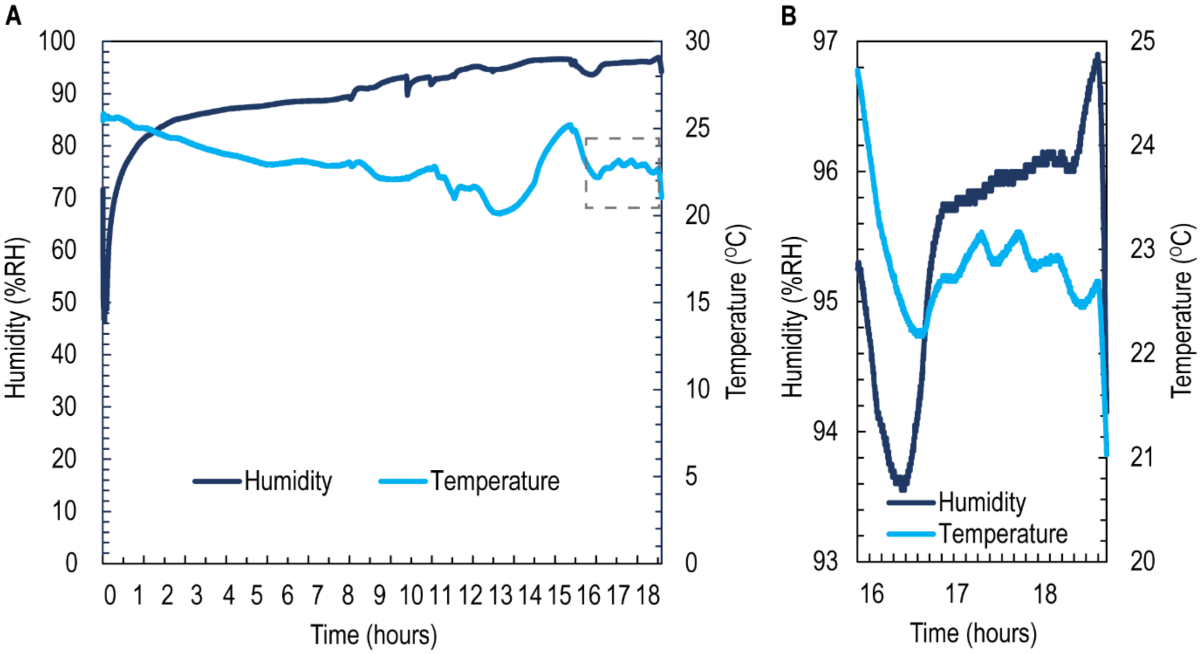


**Figure S11.** (**A**) Sample data demonstrates that the box maintains humidity levels above 90%, even after being opened four times for sample handling following the initial hydration at $t$ = 0. (**B**) Inset of (**A**) showing a more detailed view of the humidity and temperature fluctuations between $t$ = 16 h and $t$ = 19 h.

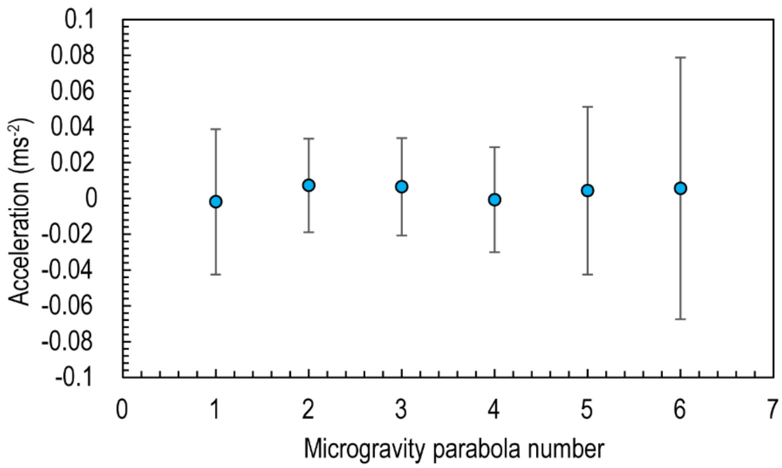


**Figure S12.** Sample data indicating that microgravity conditions were consistently achieved during each parabola of the parabolic flight.

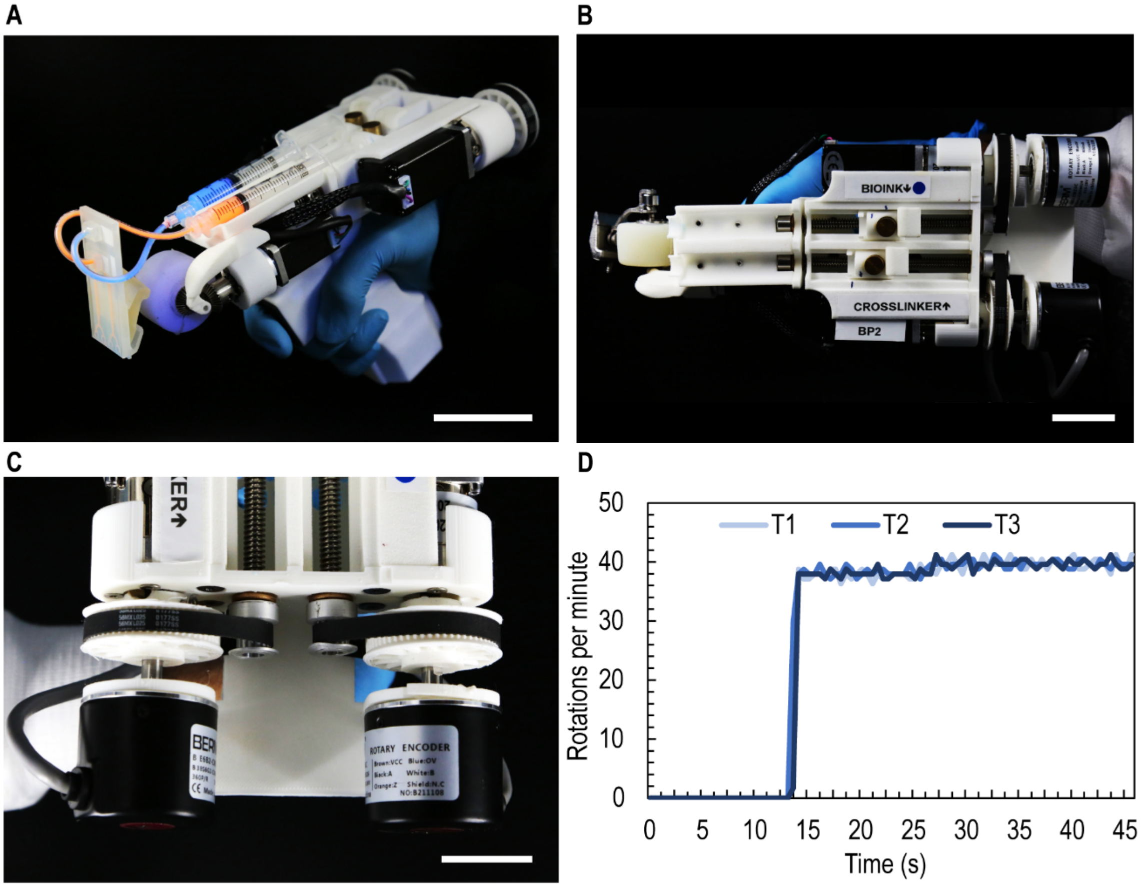


**Figure S13.** (**A**) Photograph of the handheld bioprinter, adapted from a previously published version,[S1] loaded with bi-phasic bioink (blue) and crosslinker thrombin (orange) (scale bar, 5 cm). (**B**) and (**C**) Photographs showing assembly with stepper motors fitted with rotary encoders to ensure precise control over rotational velocity (scale bars, 5 cm and 2.5 cm, respectively). (**D**) Set point for bioink delivery is 40 RPM, with motors maintaining consistent rotational speed across trials 1, 2, and 3 (T1-T3).

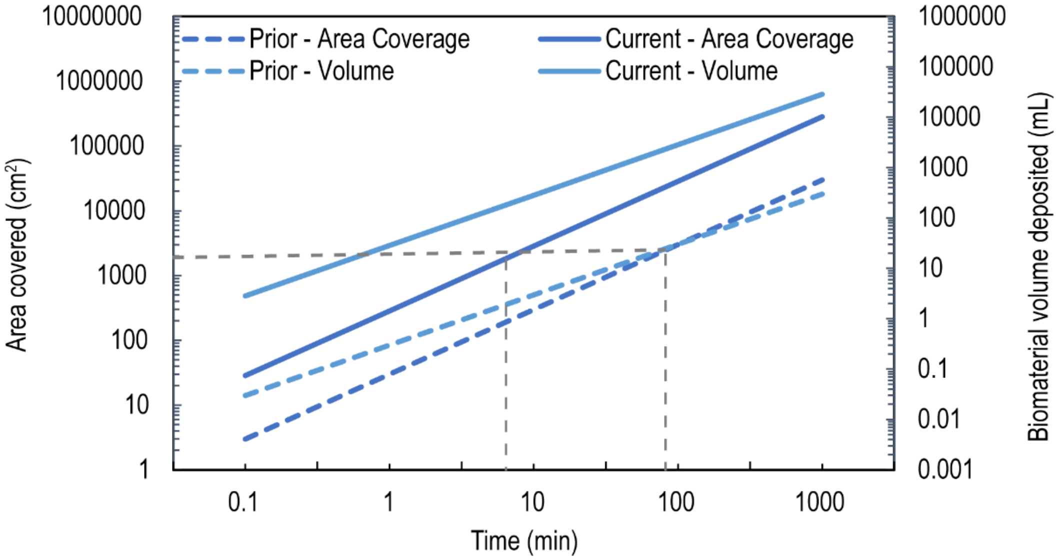


**Figure S14.** Theoretical area coverage and deposited bioink volume of the current handheld bioprinter (solid lines) are significantly higher than previously published results (dashed lines)[S1]. The gray dotted lines indicate the average burn area for human posterior trunk,[S1] with estimated printing times indicated for both the current and previously published handheld bioprinters.

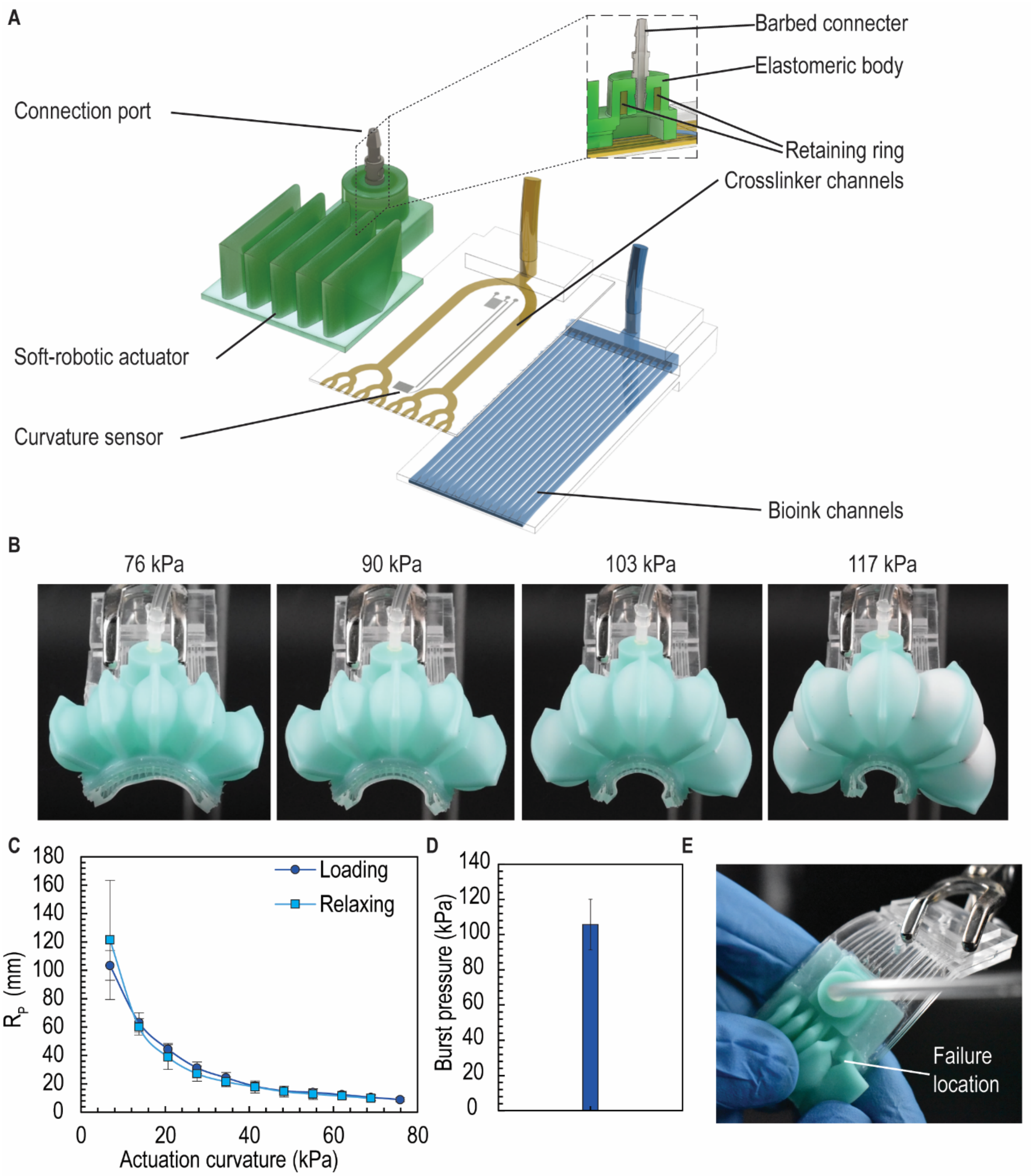


**Figure S15.** Assembly and mechanical robustness of soft robotically actuated multinozzle printhead. (**A**) Exploded view of printhead showing the microchannel layer for bi-phasic bioink delivery (bottom), cross-linker delivery and curvature sensing layer (middle) and soft robotic layer (top). (**B**) Photographs of the inflated printhead morphology near maximum actuation pressure (burst pressure). (**C**) Radius of curvature, $R_C$, of printhead measured under loading (increasing pressure) and relaxing (decreasing pressure) conditions. (**D**) Measured burst pressure of the soft robotic system. (**E**) Typical failure location observed after bursting, with the pneumatic world-to-chip interconnect remaining intact.

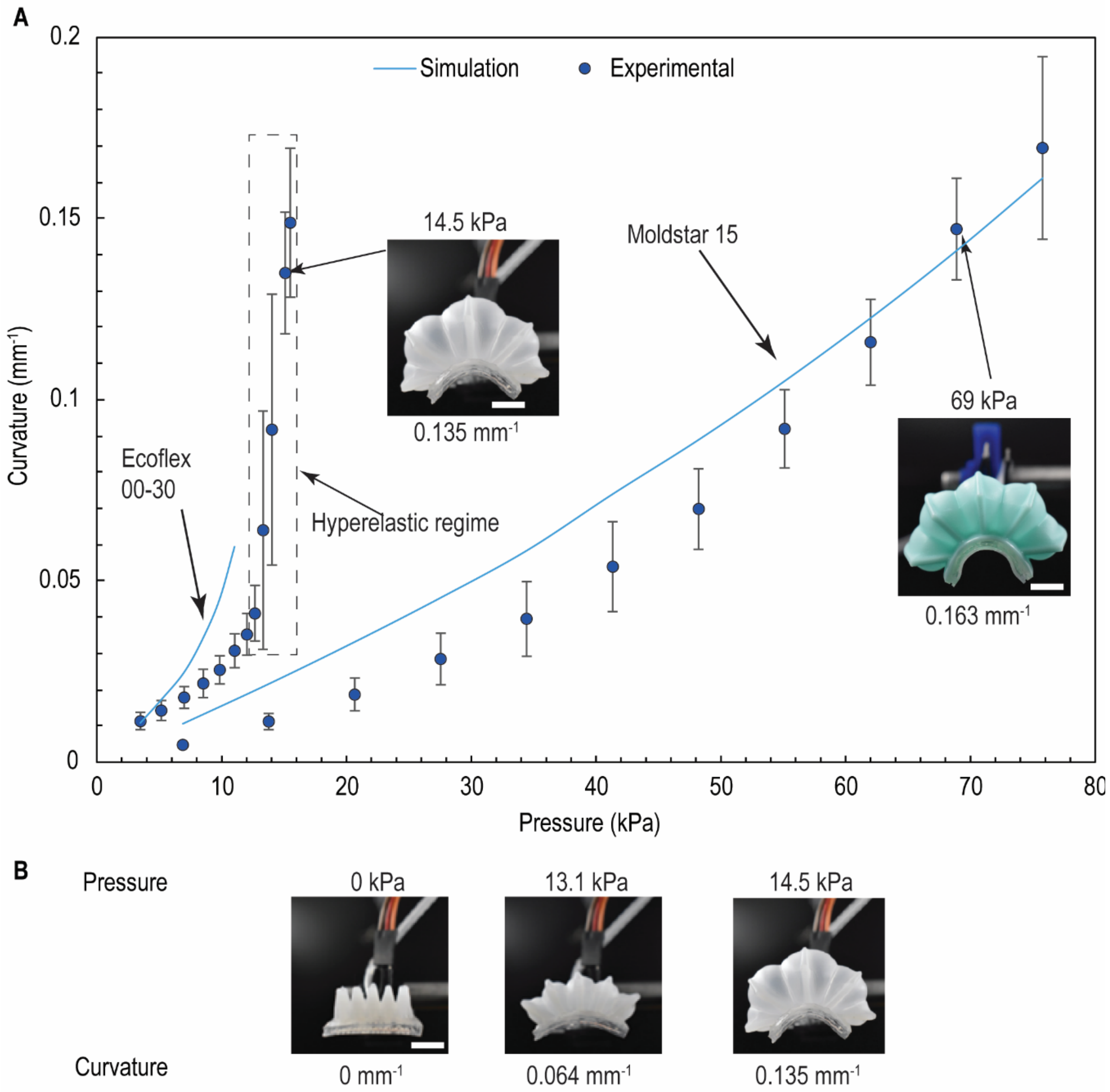


**Figure S16.** (**A**) Comparison of simulation and experimental results for soft robotically actuated printheads with actuation layers molded from two elastomers: Ecoflex 00-30 and Moldstar 15. Ecoflex 00-30 leads to undesirable hyperelastic overexpansion of the center compartments, resulting in a mismatch between the finite element simulation and experimental actuation curvature. In contrast, Moldstar 15 exhibits predictable actuation behavior and is used for subsequent experiments. (**B**) Frontal photographs showing the nonlinear change in printhead curvature and deformation for an actuation layer molded from Ecoflex 00-30 operating in the hyperelastic regime (scale bar, 10 mm).

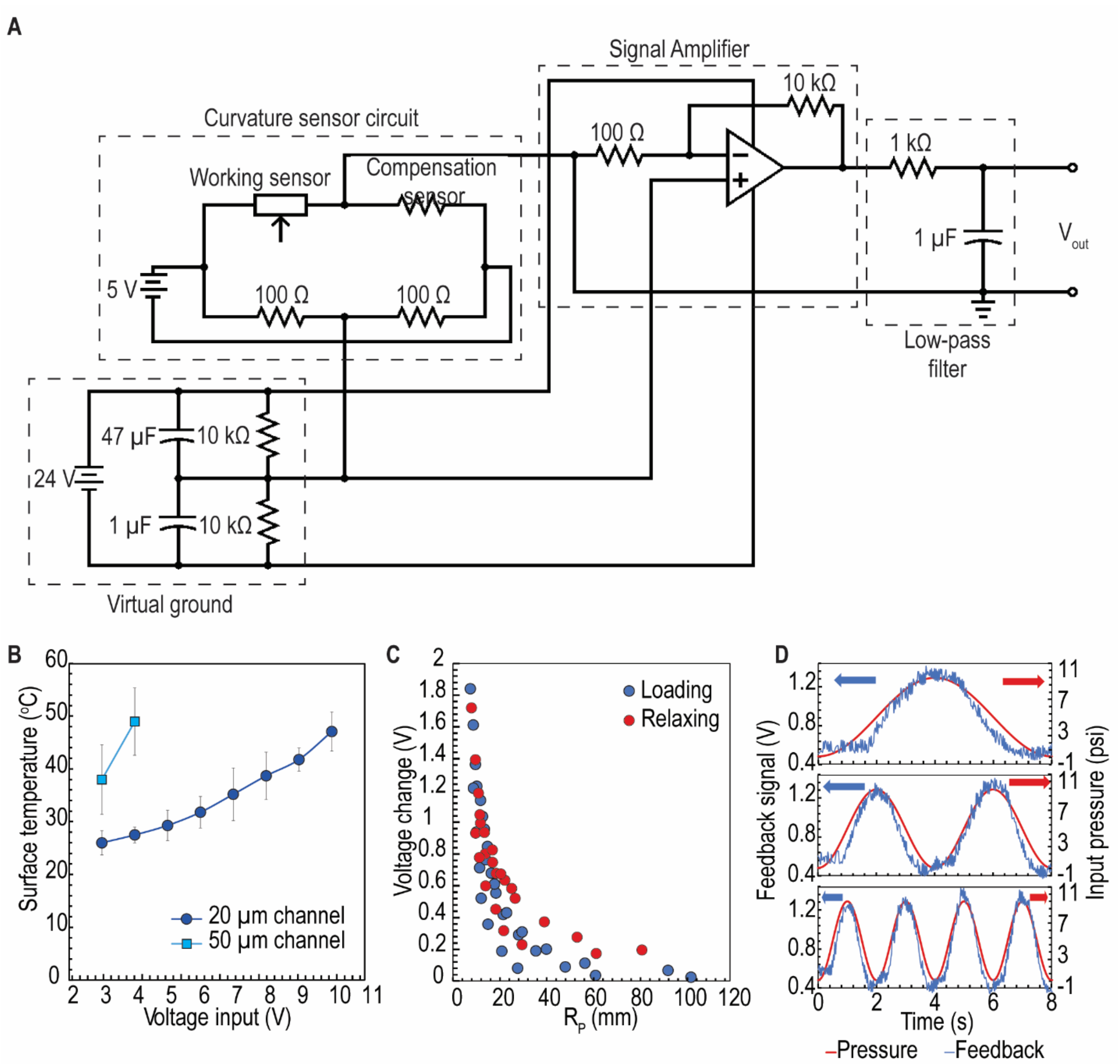


**Figure S17.** (**A**) Circuit diagram of the complete curvature sensing system. (**B**) Steady-state temperature measured at the upper surface of fluidic feature for different sensor microchannel widths. In the final sensor design, temperature remains below 30°C with a 5V input voltage. (**C**) Voltage output of the curvature sensor as a function of the printhead curvature radius ($R_P$). (**D**) Open-loop performance of the curvature sensor feedback system, demonstrating sensor feedback without active control.

A B C D E

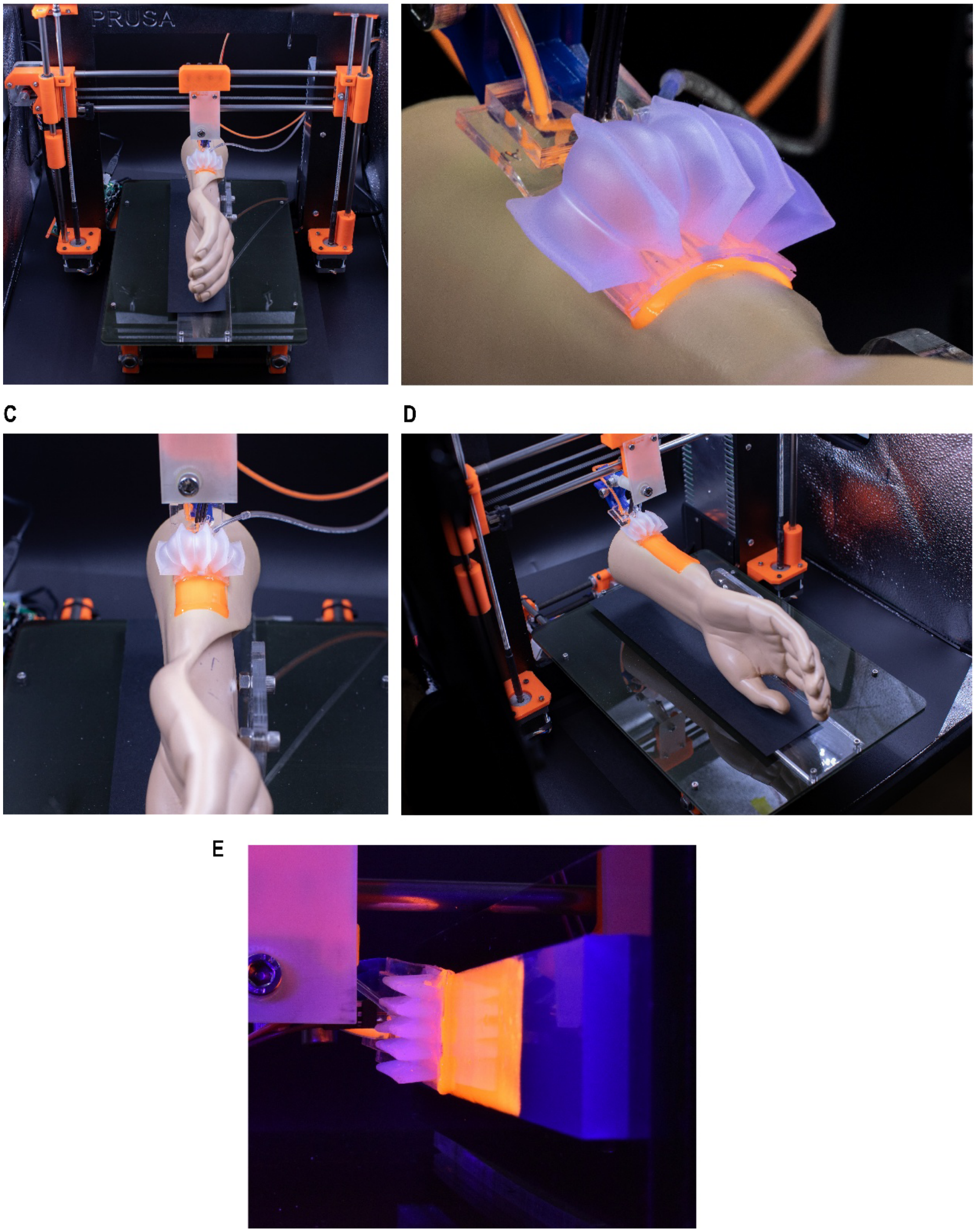


**Figure S18.** (**A**) A modified 3D printer equipped with a soft robotically actuated printhead, with the actuation layer molded from Ecoflex 00-30. The same setup was used for printheads molded with Moldstar 15. (**B-D**) During operation, the printhead is pressurized while biphasic bioink and cross-linker are separately delivered via syringe pumps. The printhead is linearly translated to deposit granular bioink onto a stationary phantom arm segment with a target curvature radius ($R_2$) of 25 mm transverse to the printing direction. (**E**) The printhead enables deposition of the bi-phasic bioinks on vertical surfaces.

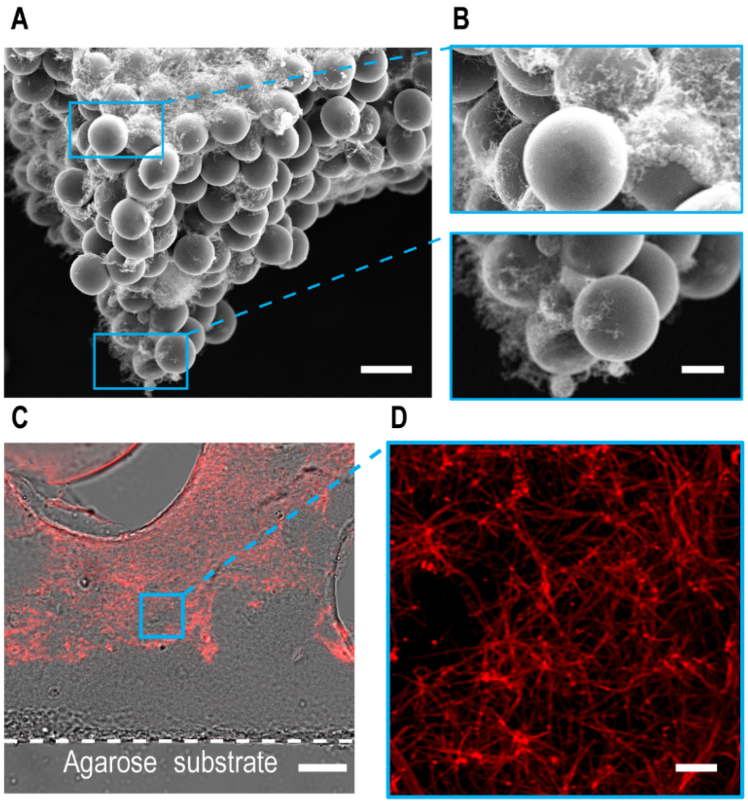


**Figure S19. (A)** Scanning electron micrograph of the cross-section of a printed sheet at an inclined orientation (45°), confirming fibrin clot formation (scale bar, 50 μm). **(B)** Higher magnification images showing the top (i) and the bottom surfaces of the sheet (ii) (scale bar, i. and ii. 20 μm). (C) Anti-fibrinogen staining confirms clot formation at the bottom of the printed sheets, revealing **(D)** the expected fibrin fibers (scale bar, **(C)** 20 μm and **(D)** 5 μm).

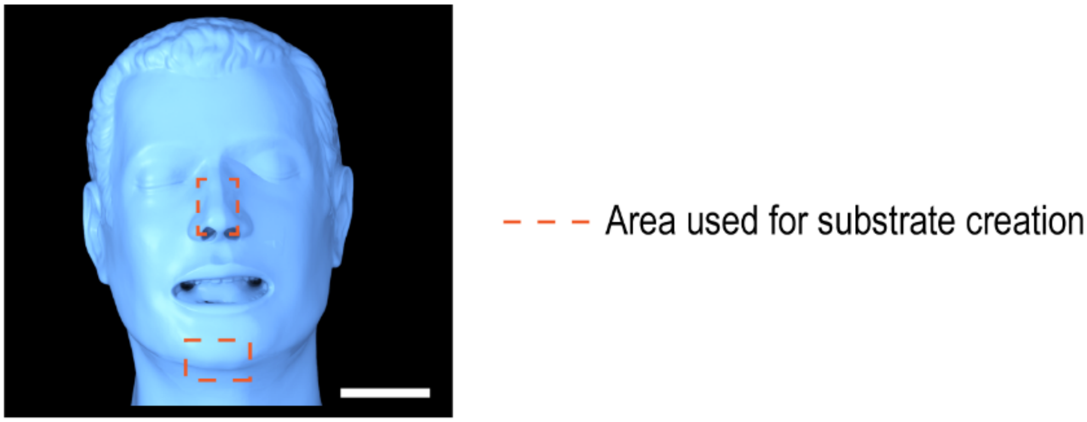


**Figure S20.** 3D laser scan of the phantom model used to create physiologically relevant substrates (scale bar, 6 cm). Regions selected for substrate fabrication are outlined with dotted lines.

**Table S1.**
List of mean and Gaussian curvature of all surface profiles used in this study. For physiological facial features, the values are comparable to previous study on human facial curvatures.

| SURFACE FEATURES | MEAN CURVATURE ($MM^{-1}$) | PREVIOUS STUDY | GAUSSIAN CURVATURE ($MM^{-2}$) | PREVIOUS STUDY |
|---|---|---|---|---|
| **CHIN** | 0.047 ± 0.002 | 0.03 - 0.04 | 0.0014 ± 0.0002 | 0.0009 - 0.0013 |
| **NOSE** | 0.082 ± 0.021 | 0.045 - 0.085 | 0.0042 ± 0.0050 | -0.002 - 0.007 |
| **LINEAR CHANGE** | 0 - 0.05 | N/A | 0 | N/A |
| **SINUSOIDAL CHANGE** | 0.02 - 0.05 | N/A | 0 | N/A |
